# RecN and RecA orchestrate an ordered DNA supercompaction response following ciprofloxacin exposure in *Escherichia coli*

**DOI:** 10.1101/2024.11.15.623168

**Authors:** Krister Vikedal, Synnøve Brandt Ræder, Ida Mathilde Riisnæs, Magnar Bjørås, James Booth, Kirsten Skarstad, Emily Helgesen

## Abstract

Fluoroquinolones induce double-strand breaks in bacterial DNA, triggering the SOS response, a major DNA damage response that ensures the expression of repair proteins but also promotes the emergence and spread of antibiotic resistance. Fluoroquinolone resistance, particularly in *Escherichia coli*, is a growing global health concern. Understanding bacterial responses to these antibiotics is critical for developing preventive strategies and novel treatments to combat resistance development. This study investigates DNA morphology in *E. coli* following exposure to ciprofloxacin, a fluoroquinolone antibiotic. We show that ciprofloxacin induces a stepwise DNA reorganization, culminating in a highly dense nucleoid structure at midcell — a process we term DNA supercompaction. Live cell imaging revealed that RecN, a structural maintenance of chromosomes (SMC)-like protein, is required for DNA supercompaction, and that RecN’s dynamics and activity in this response depend on RecA. Additionally, RecN and RecA frequently colocalized at nucleoid-associated positions. We suggest that RecN and RecA play active roles in DNA supercompaction following severe DNA damage, that their interplay is part of a prompt universal survival response to DNA double-strand breaks in *E. coli,* and that the extent of the compaction response depends on the number of double-strand breaks.

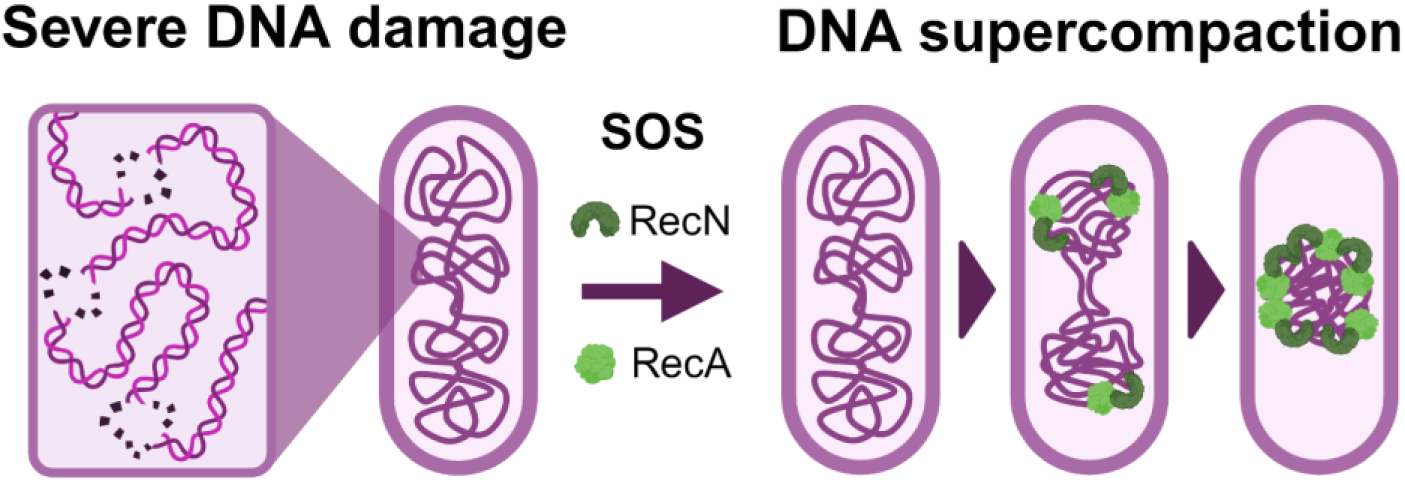
GRAPHICAL ABSTRACT.

## INTRODUCTION

Resistance to fluoroquinolone-class antibiotics poses a growing global health concern (1). In 2019, fluoroquinolone resistance was responsible for approximately a quarter of worldwide deaths attributable to antimicrobial resistance, with most of these deaths linked to fluoroquinolone-resistant *Escherichia coli* (1). These antibiotics, such as ciprofloxacin (CIP), target and inhibit DNA gyrase and topoisomerase IV, enzymes critical for maintenance of DNA supercoiling and important for DNA replication and repair. Inhibition of these enzymes by CIP leads to DNA double-strand breaks (DSBs), ultimately resulting in cell death (2, 3).

*E. coli* respond to DSBs through highly efficient, conserved mechanisms to preserve chromosomal integrity and ensure survival (4). A major repair pathway is homologous recombination, where intact double-stranded DNA (dsDNA) serves as a template to repair broken DNA (5, 6). RecBCD and RecA enzymes play pivotal roles in this process (7). RecBCD processes DSB ends to create single-stranded DNA (ssDNA) (8, 9). Upon recognition of the Chi sequence by RecBCD, RecA is loaded onto these ssDNA regions, forming RecA-ssDNA filaments that ultimately facilitate the search for and invasion of homologous dsDNA to mediate genetic exchange and repair (6, 10, 11).

Besides its central role in homologous recombination, RecA is also critical for the induction of the SOS response, the major DNA damage response. RecA-ssDNA filaments promote the autocatalytic degradation of the SOS response repressor LexA, triggering the expression of genes needed for DSB repair and survival (12–14). The SOS response not only governs DNA repair mechanisms but also contributes to the emergence and spread of antibiotic resistance (15, 16). Therefore, understanding the facets of the SOS response is critical for developing strategies to manage increasing antibiotic resistance.

DNA damage caused by genotoxic agents such as mitomycin C (MMC), nalidixic acid, or UV irradiation induces significant morphological changes in the bacterial nucleoid (17–21). These agents have been associated with varying DNA compaction phenotypes and kinetics. For instance, nalidixic acid exposure has been observed to result in a dense, persistent midcell nucleoid (17), whereas UV irradiation can lead to transient DNA compaction at both quarter and midcell positions followed by prolonged decompaction (18). Similarly, MMC exposure has been reported to trigger transient compaction followed by decompaction (19). Notably, RecA is necessary for DNA compaction induced by UV and MMC but has been reported not to be required for the initial stages of compaction induced by nalidixic acid (17–19). Another critical player in the DNA damage response is the structural maintenance of chromosomes (SMC)-like protein RecN, which influences both DNA compaction and repair after exposure to genotoxic agents (18, 21). It remains to be determined whether the various DNA compaction phenomena aid DNA repair or are instead consequences of macromolecular crowding and other intracellular changes related to unknown survival mechanisms.

RecN, a highly conserved protein, is one of the most abundantly produced proteins during the SOS response (13, 22, 23). The *recN* gene is rapidly expressed in response to DNA damage (13, 24) but is also rapidly digested by the ClpXP protease system to ensure high levels only upon DNA damage (25, 26). RecN has been suggested to play a crucial role in aiding DSB repair, and mutants lacking RecN exhibit heightened sensitivity to ionizing radiation, MMC, and I-SceI induced DSBs (27–29).

The exact mechanism by which RecN aids DSB repair is not fully understood. In vitro studies have revealed that RecN binds preferentially to RecA-bound ssDNA at DSBs (30), enabling it to bridge a dsDNA molecule via ATP-dependent topological entrapment (23, 30–32). In vivo, RecN has been shown to promote contacts between newly replicated homologous DNA strands after MMC exposure (19). GFP-tagged RecN localizes to the nucleoid following DNA damage (26), and colocalization with RecA at nucleoid gaps upon MMC exposure has been observed (33). Additionally, RecA is necessary for RecN nucleoid localization, and RecA repair activities appear to depend on RecN (33, 34). Importantly, cells lacking RecN are unable to perform DNA compaction effectively after exposure to various genotoxic agents (18, 21).

In this study, we investigate the change in DNA morphology following CIP exposure in *E. coli.* Our findings reveal that CIP exposure leads to a stepwise reorganization of DNA towards midcell — a process we term DNA supercompaction. We also find that the process requires both RecN and RecA, in what appears to be an active interplay. We suggest that DNA supercompaction is part of a prompt universal survival response to severe DNA damage.

## MATERIAL AND METHODS

### Strain construction

Strains and plasmids used in this study are listed in Table 1 and Table 2, respectively. Most strains are derived from the *E. coli* K-12 strain BW25113, which serves as the background strain of the Keio collection (35). To examine the effect of *recN* deletion, we utilized the Keio collection strain JW5416. Standard P1 transduction procedures (36) were used to knock out *recA* and introduce *hupA100*::mCherry (generous gift from Steven Sandler (37)) for visualization of DNA in the relevant backgrounds (see Table 1). FLP recombinase (pCP20) (38) was used to remove the *kan*-cassette when both donor and recipient strains carried kanamycin resistance.

**Table 1.** *E. coli* strains used in this study. Relevant antibiotic resistances are shown in brackets. P1 transduction is shown as: P1 Donor x Recipient. *, The *lexA41* (*lexA3*) mutations correspond to the point mutations *lexA*_G85D, A132T_, as determined from sequencing data of the DM936 and KP84 strains. PMGR, Plasmid-mediated gene replacement; Kan^R^, Kanamycin resistance; Cam^R^, Chloramphenicol resistance; Amp^R^, Ampicillin resistance; Tet^R^, Tetracycline resistance; Str ^R^, Streptomycin resistance.

| Strain | Relevant genotype | Source |
| --- | --- | --- |
| BW25113 | Wild type | (40); CGSC7636 |
| SS6282 | MG1655 *hupA100::mCherry::kan*<br>*ΔattB::psulA-gfp* [Kan<sup>R</sup>] | (41); S. Sandler |
| KV21 | BW25113 *hupA100::mCherry* | This work; P1 SS6282 x<br>BW25113, kan removed with<br>pCP20 |
| JW5416 | BW25113 *ΔrecN::kan* [Kan<sup>R</sup>] | (35); Keio collection |
| MR09 | BW25113 *ΔrecN* | This work; kan removed from<br>JW5416 with pCP20 |
| MR16 | BW25113 *ΔrecN hupA100::mCherry::kan*<br>[Kan<sup>R</sup>] | This work; P1 SS6282 x MR09 |
| KV60 | KV21 + pSG101 (pSOS) [Amp<sup>R</sup>] | This work |
| KV61 | KV21 + pTF271 (pARA) [Amp<sup>R</sup>, Cam<sup>R</sup>] | This work |
| KV62 | MR16 + pSG101 (pSOS) [Kan<sup>R</sup>, Amp<sup>R</sup>] | This work |
| KV63 | MR16 + pTF271 (pARA) [Kan<sup>R</sup>, Amp<sup>R</sup>,<br>Cam<sup>R</sup>] | This work |
| JW2669 | BW25113 *ΔrecA::kan* [Kan<sup>R</sup>] | (35); Keio collection |
| KV66 | KV61 *ΔrecA::kan* + pSG101 (pSOS) [Kan<sup>R</sup>,<br>Amp<sup>R</sup>] | This work; P1 JW2669 x KV60 |
| KV67 | KV61 *ΔrecA::kan* + pTF271 (pARA) [Kan<sup>R</sup>,<br>Amp<sup>R</sup>, Cam<sup>R</sup>] | This work; P1 JW2669 x KV61 |
| KV68 | BW25113 *ΔrecN hupA100::mCherry* | This work; kan removed from<br>MR16 with pCP20 |
| KV69 | BW25113 *ΔrecN ΔrecA::kan*<br>*hupA100::mCherry* [Kan<sup>R</sup>] | This work; P1 JW2669 x KV68 |
| KV70 | KV69 + pSG101 (pSOS) [Kan<sup>R</sup>, Amp<sup>R</sup>] | This work |
| KV71 | KV69 + pTF271 (pARA) [Kan<sup>R</sup>, Amp<sup>R</sup>, Cam<sup>R</sup>] | This work |
| KV75 | BW25113 *recA::mCherry* [Cam<sup>R</sup>] | This work; PMGR in BW25113<br>with pDL5196 |
| KV76 | JW5416 *recA::mCherry* [Kan<sup>R</sup>, Cam<sup>R</sup>] | This work; PMGR in JW5416<br>with pDL5196 |
| KV77 | KV75 + pSG101 (pSOS) [Cam<sup>R</sup>, Amp<sup>R</sup>] | This work |
| KV78 | KV76 + pSG101 (pSOS) [Kan<sup>R</sup>, Cam<sup>R</sup>, Amp<sup>R</sup>] | This work |
| KV79 | KV75 + pTF271 (pARA) [Cam<sup>R</sup>, Amp<sup>R</sup>] | This work |
| KV80 | KV76 + pTF271 (pARA) [Kan<sup>R</sup>, Cam<sup>R</sup>, Amp<sup>R</sup>] | This work |
| DM936* | *lexA41 (lexA3) recA1* | (42, 43); Lab collection |
| KP84* | *lexA41 (lexA3) [Tet<sup>R</sup>]* | (44); Lab collection |
| BTH101 | *cya-99 [Str<sup>R</sup>]* | Euromedex (Cat. no. EUK001) |

**Table 2.**
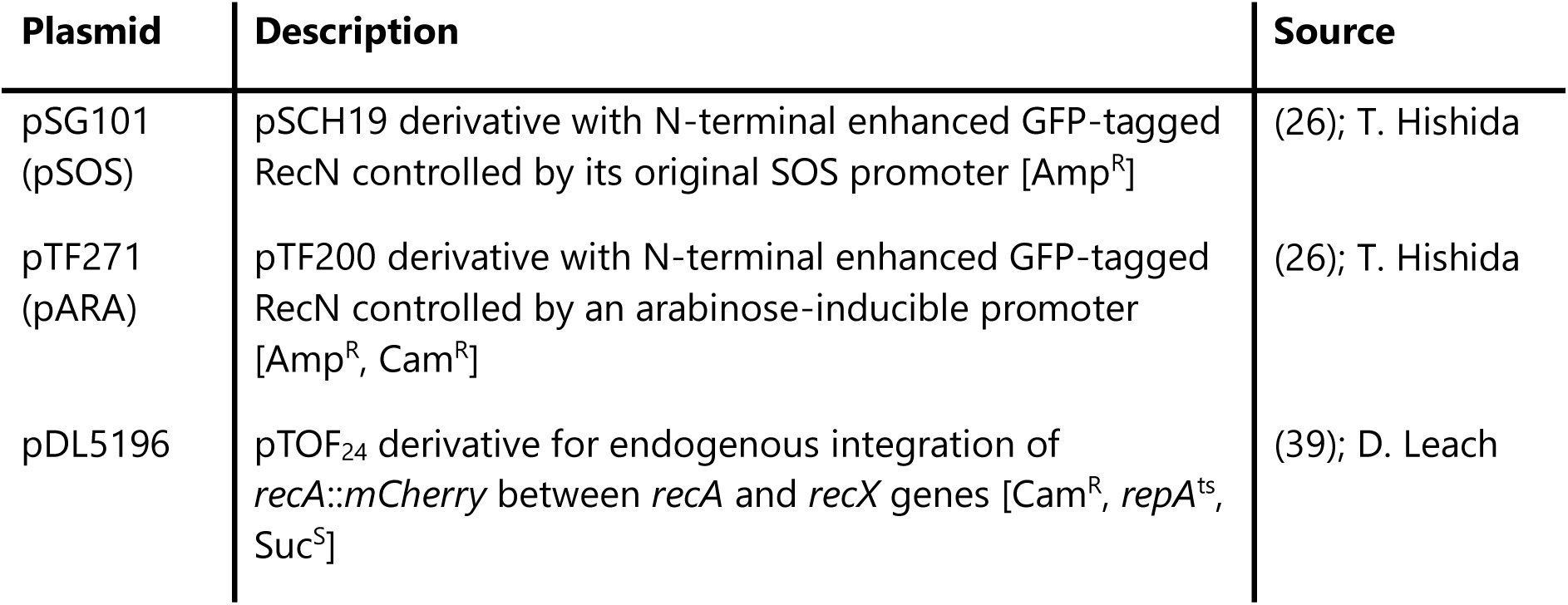
Plasmids used for main experiments in this study. Cam^R^ = Chloramphenicol resistance; Amp^R^ = Ampicillin resistance; *repA*^ts^ = Temperature-sensitive origin of replication (replicates at 30°C); Suc^S^ = Sucrose sensitive (5% wt/vol).

For studying the localization and dynamics of RecN, we employed plasmids encoding GFP-RecN (generous gifts from Takashi Hishida (26)), regulated either by the native SOS promoter (pSOS) or an arabinose-inducible promoter (pARA). The plasmids were purified using a Miniprep kit (Qiagen, cat. no. 27104) and introduced into relevant backgrounds using electroporation transformation (see details below).

To visualize RecA, *recA-mCherry* was used, chromosomally expressed in tandem with the endogenous *recA* gene. This construct was introduced by plasmid-mediated gene replacement (see details below), following electroporation transformation of the pDL5196 plasmid (generous gift from David Leach (39)) into relevant backgrounds.

### Growth conditions

Strains were streaked directly from frozen stocks onto LB-agar plates and incubated overnight. The next day, 5-10 colonies of each strain were picked and incubated in LB medium overnight with shaking. On the day of the experiment, the overnight cultures (ONCs) were diluted 1:200 into fresh LB medium and cultured at 37°C to exponential phase (OD_600_ around 0.4) before treatment or fixation unless otherwise specified. For strains carrying the pARA plasmid, arabinose (0.05% final concentration) was added when OD_600_ reached 0.15 and the cultures were incubated for another 60 minutes before CIP treatment. A dose of 10 µg/mL CIP was used in this study, except in specified cases where the minimum inhibitory concentration (MIC) of 20 ng/mL was used. Antibiotics were used as indicated in Table 1 and 2 with the following concentrations: 30 µg/mL Kanamycin, 100 µg/mL Ampicillin, 20 µg/mL Chloramphenicol, 5 µg/mL Tetracycline, and 100 µg/mL Streptomycin.

### Electroporation transformation

Electroporation was used to transform plasmids into recipient strains. The recipient strains were first cultured to exponential phase and then rendered electrocompetent through two rounds of washing and resuspension in MQ water with 10% glycerol while kept on ice. For electroporation, 50 µL of electrocompetent cells and 50-100 ng of purified plasmid were added to an electroporation cuvette with a 1 mm gap (Thermo Fisher, P41050), and an exponential pulse of 1.35 kV, 600 Ω, and 10 µF was applied using a Gene Pulser II (Bio-Rad).

Immediately after electroporation, the cells were suspended in 700 µL SOC medium and incubated at 37°C with shaking for 1-2 hours for recovery. Then, 100 µL of the recovered cell culture was plated on LB-agar with appropriate antibiotics and incubated at 37°C overnight. The next day, single transformant colonies were picked and grown to saturation before preparation of frozen stocks.

### Plasmid-mediated gene replacement

A protocol from David Leach’s lab (39), based on work by Link et al. (45) and Merlin et al. (46), was used for integration of the *recA-mCherry* construct between the chromosomal *recA* and *recX* genes using the pDL5196 plasmid. This plasmid carries *recA-mCherry,* chloramphenicol resistance (Cam^R^), a temperature-sensitive origin of replication (*repA*^ts^), and sucrose sensitivity (*sacB*) (see Table 2).

The pDL5196 plasmid was introduced into the strain of interest via electroporation transformation, recovering cells at 30°C before they were plated on LB plates containing chloramphenicol for selection. To eliminate the pDL5196 plasmid and select for cells that had integrated recA-mCherry through recombination, colonies were restreaked onto chloramphenicol plates and incubated at 42°C overnight. This selection process was repeated the next day. To confirm plasmid loss, individual colonies were cultured in LB without chloramphenicol overnight and screened on LB-sucrose plates. Finally, the presence of mCherry fluorescence was verified by fluorescence microscopy.

### Live-cell imaging with microfluidic setup

To examine the cells’ immediate response to CIP exposure, we employed a microfluidic setup to image cells using a Zeiss Axio Observer Z1 widefield inverted microscope with a 63x oil objective (Zeiss Plan Apochromat 1.4 NA, DIC), a Colibri 7 LED light source, a Hamamatsu ORCA-Flash4.0 V3 digital CMOS camera, as well as a heated incubation chamber and mounting frame, both maintained at 37°C.

Cells were cultured to exponential phase and immobilized on a chitosan-coated coverslip (Chitozen, Idylle Labs, TMI-CHI-7525) within microfluidic channels (Ibidi, 80608) following Idylle Labs’ standard protocol. Pre-cleaned microfluidic tubes were connected to the inlet and outlet of the microfluidic channel to enable flow of growth medium. LB½ medium (LB diluted 1:1 with Milli-Q water) was used to reduce background noise during imaging, and care was taken to not introduce air bubbles during setup. A peristaltic pump (Ismatec, ISM930C) was used to apply medium flow through the channel at 2 mL/min for 5 minutes to clear non-attached cells.

Cells in the microfluidic channels were imaged at 2-minute intervals and allowed to grow unchallenged for 30 minutes to verify normal cell growth and division. Next, medium with CIP (10 µg/mL) was introduced by applying flow for 10 minutes to ensure complete medium exchange. To account for focus drift, Z-stacks of 7 slices at 0.50 µm intervals with manually adjusted midpoints were captured. Transmitted light (TL) and mCherry fluorescence channels were imaged using 2×2 binning. The mCherry was excited at 540-570 nm and emissions collected at 570-640 nm. Because of signs of bleaching or photodamage, we excluded frames after 60 minutes of timelapse imaging from the presented figures.

### Live-cell spinning disk microscopy

To explore details of DNA supercompaction and the interplay between RecN and RecA, we employed spinning disk microscopy with improved focus stability and reduced photodamage effects. The Nikon Eclipse Ti2-E inverted microscope was equipped with a 60x oil objective (Nikon Plan Apochromat λD 60x 1.42 NA, DIC), a CrestOptics X-Light V3 spinning disk confocal module (50:400 µm spinning disk), a Lumencor Celeste multi-line laser, two Teledyne Photometrics Kinetix sCMOS cameras, as well as a stage-top incubator chamber and an objective heater, both maintained at 37°C. The microscope’s Perfect Focus System (automatic focus), guided by near-infrared light, ensured that cells remained consistently in focus without requiring the use of Z-stacks.

Cells were cultured to exponential phase, as previously described. Immediately after addition of CIP, cultures were gently shaken for 1 minute before 10 µL was transferred to an LB½-agar pad (1% agarose in LB½ medium) pre-made on a microscope glass slide within a Gene Frame (Thermo Scientific, no. AB0576). Once dried, the agar pad was sealed with a cover slip (#1.5 thickness). Imaging commenced 10 minutes after CIP exposure, with acquisition of three locations at either 10-second or 2-minute intervals, as specified in each case.

Three channels were used for imaging: a TL channel for cell outlines and fluorescence channels for GFP and mCherry. GFP was excited at 477 nm and its emissions collected at 501-521 nm, whereas mCherry was excited at 546 nm and its emissions collected at 580-610 nm. We did not observe bleaching or photodamage throughout our timelapse experiments.

### Microscopy of DNA in fixed cells

The DNA compaction phenotype of strains without chromosomally expressed HU-mCherry was investigated before and after CIP exposure using conventional DNA staining with Hoechst 33258. Cells were grown to exponential phase in LB and fixed with ethanol (50% final concentration), before storage at 4°C until further use. Prior to imaging, samples were stained with 5 µg/mL Hoechst 33258 in PBS for 10 minutes, washed with PBS, and up-concentrated 2-4 times.

#### Imaging of strains with temperature-regulated SOS induction to evaluate the role of RecA in DNA supercompaction

DM936 and KP84 strains were cultured at 30°C, 37°C, or 42°C to achieve varying levels of SOS induction. Samples were fixed prior to treatment and after 20, 40, and 60 minutes of CIP exposure. After staining with Hoechst 33258, samples were prepared for imaging in a glass-bottom 384-well plate (Fisher Scientific, no. 10687354), pre-coated with Poly-L-Lysine (Merck Life Science, no. P8920). The plate was centrifuged for 10 minutes at 700 x *g* to ensure cell attachment.

For imaging, an ImageXpress Micro Confocal High-Content Imaging System (Molecular Devices) was used, with a 60x air objective (Nikon CFI Plan Apochromat Lambda 60XC, 0.95 NA) and an Andor Zyla sCMOS camera. Imaging and focusing were fully automatic. Four locations, evenly spaced by 600 µm, were imaged per well using two channels: a TL channel for detecting cell outlines, and a fluorescence channel for Hoechst 33258 with excitation at 350-404 nm and emission collection at 417-477 nm.

#### Imaging of DNA in recA-mCherry strains

KV75, KV76, KV77, KV78, KV79, and KV80 strains, all containing the *recA-mCherry* construct, were cultured at 37°C, with samples fixed prior to treatment and after 15 and 30 minutes of CIP exposure. After Hoechst staining, 10 µL of sample was added to an agar pad on a glass slide pre-made within a Gene Frame (Thermo Scientific, no. AB0576), sealed with a cover slip (#1.5 thickness) once dried.

We used a Leica DM6000 B microscope with a 100x oil objective (Leica HCX Plan Apochromat 100x 1.40 NA, PH3 CS), a Leica EL6000 metal-halide light source, and a Hamamatsu C9100-14 EM-CCD camera. Three locations across the agar pad were imaged to include many cells in the analysis. Each image included a phase-contrast channel for cell outlines, and a fluorescence channel for Hoechst 33258 with excitation at 334-406 nm and emission collection at 407-497 nm.

### Image processing and analysis

To process and analyze images, we used the open-source software Fiji (ImageJ) (47) and its plugins Coli-Inspector (48) and MicrobeJ (49). Coli-Inspector was used to analyze images from the live-cell microfluidic setup, and from the fixed cells with temperature-regulated SOS induction (DM936 and KP84). MicrobeJ was used to analyze all remaining images. All scripts and templates used for processing and analysis in this study are available in a Zenodo repository (see Data Availability).

All images were preprocessed to remove background noise, with methods depending on the microscopy setup used. For images from the microfluidic setup, only the best-focused slice from the Z-stack was selected for each time point. For images from spinning disk microscopy, flat-field correction was applied to all image channels, in addition to a Gaussian blur (1-2 pixels radius) for the TL channel, and a background subtraction with 1-µm sliding paraboloid for fluorescence channels. For the remaining imaging techniques, noise was removed from the TL channel by subtracting a 2-µm median-filtered version and from the fluorescence channels by a 1-µm sliding paraboloid background subtraction.

We customized the script of the ObjectJ-based Coli-Inspector plugin (version 03f) (48) to allow grouping of cells by image number, and thus by time frame (see Data Availability). Cells were first segmented using the default method with adjusted limitations on cell area, cell width, and circularity. We measured average symmetrical fluorescence intensity profiles along cells’ long axes for each frame and transferred these measurements to GraphPad Prism (version 10.2.0). The outer bounds of the average fluorescence peaks at 80% of their maximum intensity was determined for each time point, so that the distances between the outer bounds could be plotted to illustrate the distribution of DNA along the cells’ long axes.

For kymograph creation and DNA distribution analysis, MicrobeJ (main version 5.13p (1)) was used (49). Cells were segmented using the “Medial Axis” mode of detection with limitations on cell area, length, width, curvature, sinuosity, and angularity. Cells that could not be tracked for at least 10 consecutive frames were excluded. An additional step was applied to detect division sites and segment cells accordingly. Single-cell kymographs were created with the kymograph plotting tool in the MicrobeJ results interface from individually tracked and analyzed cells. Kymograph heat maps were generated from multiple analyzed cells with the ShapePlot tool. For DNA distribution analysis, we measured average fluorescence intensity profiles along cells’ long axes for each frame and transferred these measurements to GraphPad Prism (version 10.2.0) for plotting.

A beta-version of MicrobeJ (beta-version 5.13p (13)) (49) was used to track and analyze GFP-RecN and RecA-mCherry foci because of issues with particle tracking coordinates in the current main version. The cell segmentation method was the same as in the current main version. Fluorescent foci of GFP-RecN, HU-mCherry, or RecA-mCherry were detected inside segmented bacteria. To quantify RecN colocalization with DNA, GFP-RecN foci inside the regions of already detected HU-mCherry foci were identified without any margin. The same approach was used to identify RecA foci colocalized with RecN, but with a 1-pixel margin. For tracking of distinct RecN foci over time, an absolute coordinate system was used. Trajectories with lifespans below 5 frames were discarded. The relative midcell distances of foci were calculated by dividing their absolute distance to the parent bacteria midcell by the length of the parent bacteria. All results from detected and tracked foci were transferred to GraphPad Prism (version 10.2.0) for plotting.

### Western Blotting to compare RecN expression levels

ONCs were diluted 1:100 in 100 mL LB cultures and grown to exponential phase. For strains carrying pSOS (KV60 and KV62) and the control strains (KV21 and KV68), the cultures were grown until OD_600_ reached 0.4 before half the cultures were treated with CIP (10 µg/mL) for 20 minutes. For strains carrying pARA (KV61 and KV63), arabinose (0.05% final concentration) was added when the OD_600_ reached 0.2, and the cultures were incubated for an additional 60 minutes. CIP (10 µg/mL) was added to one half of the pARA cultures for the last 20 minutes of incubation. All cultures, including unchallenged halves, were pelleted 20 minutes after CIP addition. The pellets were washed once with PBS, snap-frozen with liquid nitrogen, and resuspended in 0.5 mL lysis buffer (50 mM Tris pH 7.5, 150 mM NaCl, 1 mg/mL lysozyme, 1 mM PMSF) while kept on ice. Samples were sonicated for 25 cycles (30 seconds on, 30 seconds off) using Bioruptor Pico (Diagenode). After sonication, samples were centrifuged at 20,000 x *g* for 20 minutes and the supernatant with protein extract was retained.

Protein extracts (40 µg) from each sample was run on 4-15% Mini-PROTEAN TGX Precast Gels (Bio-Rad) using NuPAGE LDS Sample Buffer (Final 1X) and 1 mM DTT. SeeBlue Plus2 Pre-stained Protein Standard (Invitrogen) and purified his-tagged RecN protein were used as markers. The proteins were transferred to a nitrocellulose membrane (0.2 µm) using Trans-Blot Turbo Mini Transfer Packs (Bio-Rad) at 25V, 1.35A for 10 minutes in a Trans-Blot Turbo Transfer System (Bio-Rad). Afterwards, the membranes were blocked in PBS-Tween with 5% skim milk (Merck) for 45 minutes, followed by overnight blocking in cell extract made from the KV68 (*ΔrecN*) strain (see details below). After blocking, the membranes were incubated 2 hours with primary anti-RecN antibody (rabbit anti-RecN polyclonal antibody, MyBiosource, cat. no. MBS7162137) diluted 1:200 in PBS-Tween with 5% skim milk, and then 45 minutes with secondary goat anti-rabbit IgG (H+L) HRP-conjugated antibody (MyBiosource, cat. no. MBS705310) diluted 1:10,000 in PBS-Tween with 5% skim milk. Finally, the membranes were incubated with SuperSignal West Pico PLUS Chemiluminescent Substrate (Thermo Scientific, cat. no. 34577) for chemiluminescent imaging. The Western blot membranes were imaged using a ChemiDoc MP Imaging System with Image Lab software (Bio-Rad).

RecN levels from Western blot images were quantified using Fiji (ImageJ) (47). The gel lane selection tool was used to select regions encompassing both the GFP-RecN (98 kDa) and RecN (61 kDa) proteins for each sample lane. The lane profiles were plotted and the measured signal from each lane was normalized to the wild-type strain after subtracting the background signal from the negative control. The final results were plotted using GraphPad Prism version 10.2.0.

#### Cell extracts for blocking of Western blot membranes

To make cell extracts from *ΔrecN* cells (KV68) for blocking of unspecific binding by the anti-recN antibody, ONCs were diluted 1:100 in 1L LB and grown to OD_600_ of 0.5 before addition of 10 µg/mL CIP. The cultures were pelleted after 20 minutes of CIP exposure, washed with PBS, and snap-frozen. Pellets were lysed by sonication in 10 mL PBS with 1 mg/mL lysozyme at amplitude 60 for 30 seconds in three cycles using a Vibracell VC601 sonicator with a 13 mm probe (Sonics & Materials). The lysed pellets were centrifuged at 20,000 x *g* for 20 minutes, and the resulting supernatants were collected. Subsequently, the supernatants were incubated at 4°C for 2 hours, after addition of 3 mL PBS-Tween with 10 mM EDTA and 1:200 anti-recN antibody (rabbit anti-RecN polyclonal antibody, MyBiosource, cat. no. MBS7162137). Finally, this solution was centrifuged at 4,000 x *g* for 15 minutes before the resulting supernatant was added to membranes as the cell extract for blocking.

### Survival assays

#### Spot assay

To examine if the presence of GFP-RecN, or total elevated RecN levels affected cell survival, we performed a spot survival assay. Cells were cultured at 37°C to an OD_600_ of approximately 0.1. The cultures were then split into two batches: one batch was incubated with arabinose (0.05% final concentration) for an additional 60 minutes, while the other batch was incubated for an additional 45 minutes without arabinose. Samples from each batch, before and after 20 minutes of CIP exposure, were serially diluted 10-fold with LB medium (1-10^-7^) in 96-well plates. From each dilution, 10 µL was spotted onto LB-agar plates with or without 0.05% arabinose and incubated overnight at 37°C. Colony growth was assessed the next day and the plates imaged using a GelDoc Go imaging system (Bio-Rad).

#### Time-dependent survival assay

A time-dependent survival assay was used to compare the survival of wild-type (BW25113) and *ΔrecN* (JW5416) strains after different CIP exposure times. ONCs of each strain were diluted 1:100 and cultured to OD_600_ around 0.3. Each culture was split into three parts in a 96-well plate: one part remained unchallenged, one was treated with the experimental dose of CIP (10 µg/mL), and one was treated with the MIC dose of CIP (20 ng/mL). A sample of 5 µL was collected from each part prior to treatment and at multiple time points from 1 to 120 minutes after CIP exposure. The unchallenged samples served as references. All samples were serially diluted 10-fold in 96-well plates, spotted onto LB-agar plates (5 µL), and incubated overnight at 37°C. The number of viable colonies were counted the following day to determine colony-forming units (CFU) per mL. Relative survival was calculated as the CFU/mL ratio between CIP-treated and unchallenged samples from three biological replicates.

### Bacterial two hybrid assay

A bacterial two hybrid assay kit (Euromedex, cat. no. EUK001) was used to investigate direct interaction between RecN and RecA (50)(51). The assay is based on reconstitution of two fragments (T18 and T25) into adenylate cyclase (CyaA) in a *ΔcyaA* strain (BTH101), which will lead to production and activity of β-galactosidase upon direct interaction of the assayed proteins. The *recN* and *recA* genes were cloned into pK(N)T25 and pUT18(C) vectors (GenScript) to create N– and C-terminal fusions with T25 and T18 fragments (Supplementary Table S1).

The assay was based on the kit protocol, as well as work by Mehla et al. (50) and Battesti and Bouveret (51). In short, BTH101 cells were co-transformed with combinations of pUT18(C) and pK(N)T25 derivatives. Transformants were plated on LB-agar with appropriate antibiotics and incubated at 30°C for 3 days, after which ONCs were inoculated in LB with 0.5 µM IPTG. The next day, ONCs were diluted 1:100 into fresh LB with 0.5 µM IPTG and grown to OD_600_ around 0.5. Cells were then pelleted, resuspended in PM2 buffer (70 mM Na_2_HPO_4_, 30 mM NaH_2_PO_4_, 1 mM MgSO_4_, and 0.2 mM MnSO_4_) containing β-mercaptoethanol (100 mM), and permeabilized with toluene (10%), and SDS (0.01%) at 37°C for 45 minutes. Afterwards, permeabilized cells were pelleted, and the supernatant transferred to 96-well polypropylene plates in three technical replicates. The supernatants were mixed with ONPG (*o*-nitrophenyl-β-D-galactoside) (0.67 mg/mL final concentration) and incubated at 30°C to produce yellow-colored *o*-nitrophenol in reaction to β-galactosidase. After 30-40 minutes, the reaction was stopped with sodium carbonate (200 mM), and β-galactosidase activity (*o*-nitrophenol) was measured through absorption at 405 nm (OD_405_) using a Victor Nivo multimode plate reader (PerkinElmer). The results were normalized to measurements from the positive (pKT25-zip + pUT18C-zip) control.

### Statistical analysis

Single representative replicates have been presented in figures detailing DNA distribution and RecN or RecA dynamics to accurately represent the kinetics in the population which may vary slightly between replicates. The remaining results are presented as mean ± error from biological replicates. The differences in relative midcell distance between RecA foci colocalizing with RecN and non-colocalizing RecA foci were presented as 95% confidence intervals (CIs) and analyzed using one sample t-tests with a hypothetical value of 0 in GraphPad Prism (version 10.2.0). We considered two-tailed *p*-values below 0.05 as significant and indicated this with asterisks: ns, non-significant; * *p* ≤ 0.05; ** *p* ≤ 0.01; *** *p* ≤ 0.001; **** *p* ≤ 0.0001.

## RESULTS

### Ciprofloxacin exposure leads to DNA supercompaction through a stepwise reorganization

The organization of bacterial chromosomes following DNA damage depends on the type of damaging agent and the severity of the damage (17–21, 52, 53). Given the widespread resistance to CIP, we investigated if and how CIP induces chromosomal reorganization in *E. coli*. For this purpose, we used live-cell imaging to monitor DNA morphology in exponentially growing cells containing HU-mCherry for DNA visualization (KV21, hereafter referred to as wild-type).

To establish a timeline of chromosomal reorganization after CIP exposure, we initially used a widefield microscope coupled with a microfluidic setup that allowed direct administration of CIP (10 µg/mL) during image acquisition (see Materials and methods). Prior to CIP exposure, most cells displayed two clearly separated nucleoids, one in each cell half, often with two distinct nucleoid lobes each (Figure 1A, 0 min). Following CIP exposure, we observed extensive chromosomal reorganization, culminating in a dense nucleoid structure at midcell (Figure 1A, 16 min). To quantify this reorganization, we analyzed average DNA distribution along the cells’ long axis throughout the timelapse and found that cells consistently reorganized their DNA within 8 to 20 minutes post-CIP exposure (Figure 1B).

**Figure 1.**
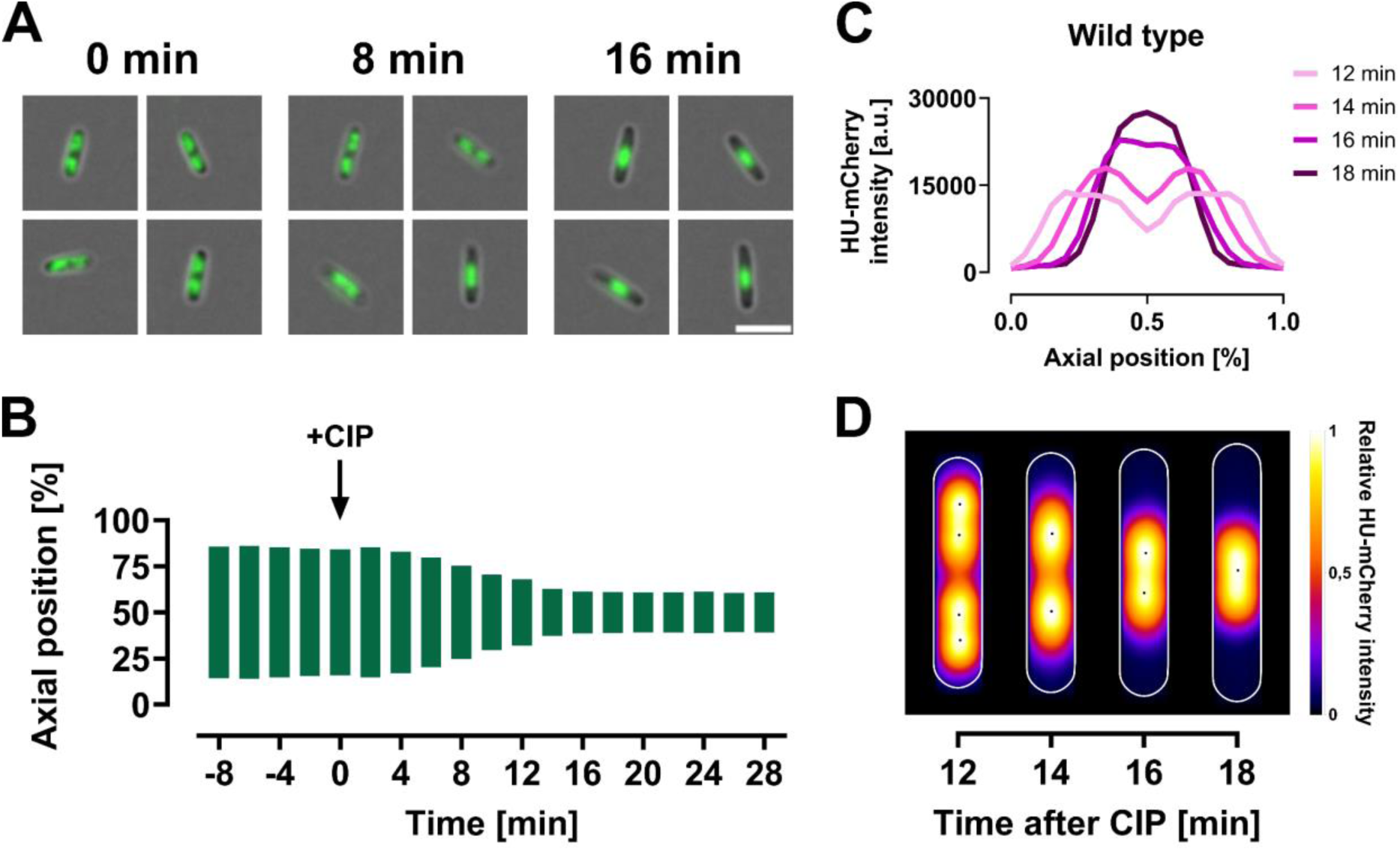
DNA supercompaction in wild-type *E. coli* (KV21) cells after CIP exposure. All cells were grown in LB at 37°C and imaged at 2-minute intervals using live-cell imaging. (**A** and **B**) Cells were immobilized in microfluidic channel slides and imaged using widefield microscopy. (**A**) Representative images of HU-mCherry labelled (green) wild-type cells at 0, 8, and 16 minutes after CIP exposure, showing DNA distribution. Scale bar: 5 µm. (**B**) Analysis of DNA distribution along the cells’ long axis before and after CIP, quantified by measuring the distance between the outer bounds of symmetrical fluorescence peaks at 80% of maximum averaged intensity for each time point. Results are averaged from 34-141 cells from a single representative biological replicate (see Materials and methods for detailed explanation). (**C** and **D**) Cells were immobilized on agar pads and imaged using spinning disk microscopy. Results shown are from images captured 12, 14, 16, and 18 minutes after CIP exposure, averaged from 40 tracked cells from a single representative biological replicate. (**C**) Average HU-mCherry fluorescence intensity along the cells’ long axis. (**D**) Kymograph heat map of relative HU-mCherry intensity distribution over time. A.u., arbitrary unit.

**Video 1.**
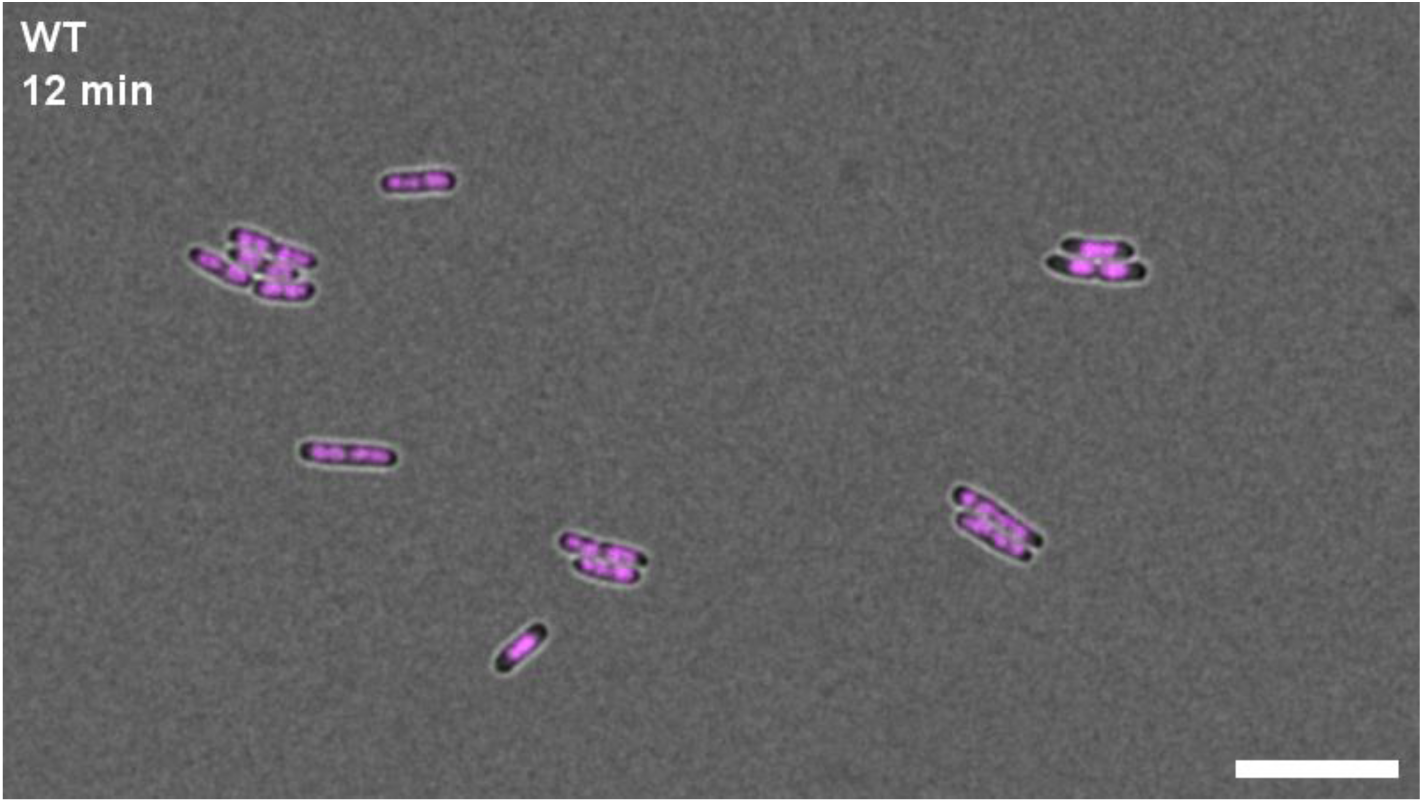
DNA supercompaction in wild-type *E. coli* (KV21) cells after CIP exposure. DNA is represented by HU-mCherry fluorescence in magenta. Cells were grown in LB at 37°C and imaged at 2-minute intervals using live-cell spinning disk microscopy, starting 12 minutes after CIP exposure. Scale bar is 10 µm. [See Video 1 in Supplementary Data]

To improve focus stability and temporal resolution without notable photobleaching, we switched to a spinning disk microscope with an automatic focus system (see Materials and methods). Using this setup, we imaged live cells at 2-minute intervals starting 10 minutes post-CIP exposure. Improved focus stability led to better separation of DNA foci represented by HU-mCherry fluorescence, allowing for identification of distinct nucleoids and nucleoid lobes. This led us to discover that the dense nucleoid structure at midcell resulted from a highly organized, stepwise compaction process and not from a sudden event. In cells with four distinct DNA foci, the two chromosomal foci within each cell half initially fused and compacted at the quarter positions (Figure 1C and D, 12-14 min). Subsequently, these two remaining DNA foci migrated toward midcell and fused (Figure 1C and D, Video 1). We term this organized process DNA supercompaction, as it appears visually distinct from the compaction caused by other DNA damaging agents such as UV (18), MMC (20) bleomycin (21), gamma-irradiation (53) or I-SceI induced damage (54).

### DNA supercompaction is dependent on RecN

We, and others, have previously demonstrated that reorganization of DNA following certain types of DNA damage largely depends on the SMC-like protein RecN (18, 19, 21). Therefore, we sought to determine if RecN is also essential for the DNA supercompaction observed after CIP exposure. We grew a *ΔrecN* strain expressing HU-mCherry (MR16) to exponential phase and imaged the cells using both widefield and spinning disk fluorescence microscopy, following the same procedures as for the wild-type cells.

In striking contrast to wild-type cells, *ΔrecN* cells exhibited minimal nucleoid dynamics after CIP exposure. The DNA foci remained nearly stationary at their pre-exposure positions, although nucleoid lobes had a slight tendency to merge at quarter positions (Figure 2A and B). Only a small fraction of *ΔrecN* cells displayed DNA foci closer to midcell (Figure 2A, 16 min). However, the DNA in these cells could not be classified as supercompact, as it occupied larger regions than the midcell DNA structure in wild-type cells after CIP exposure.

**Figure 2.**
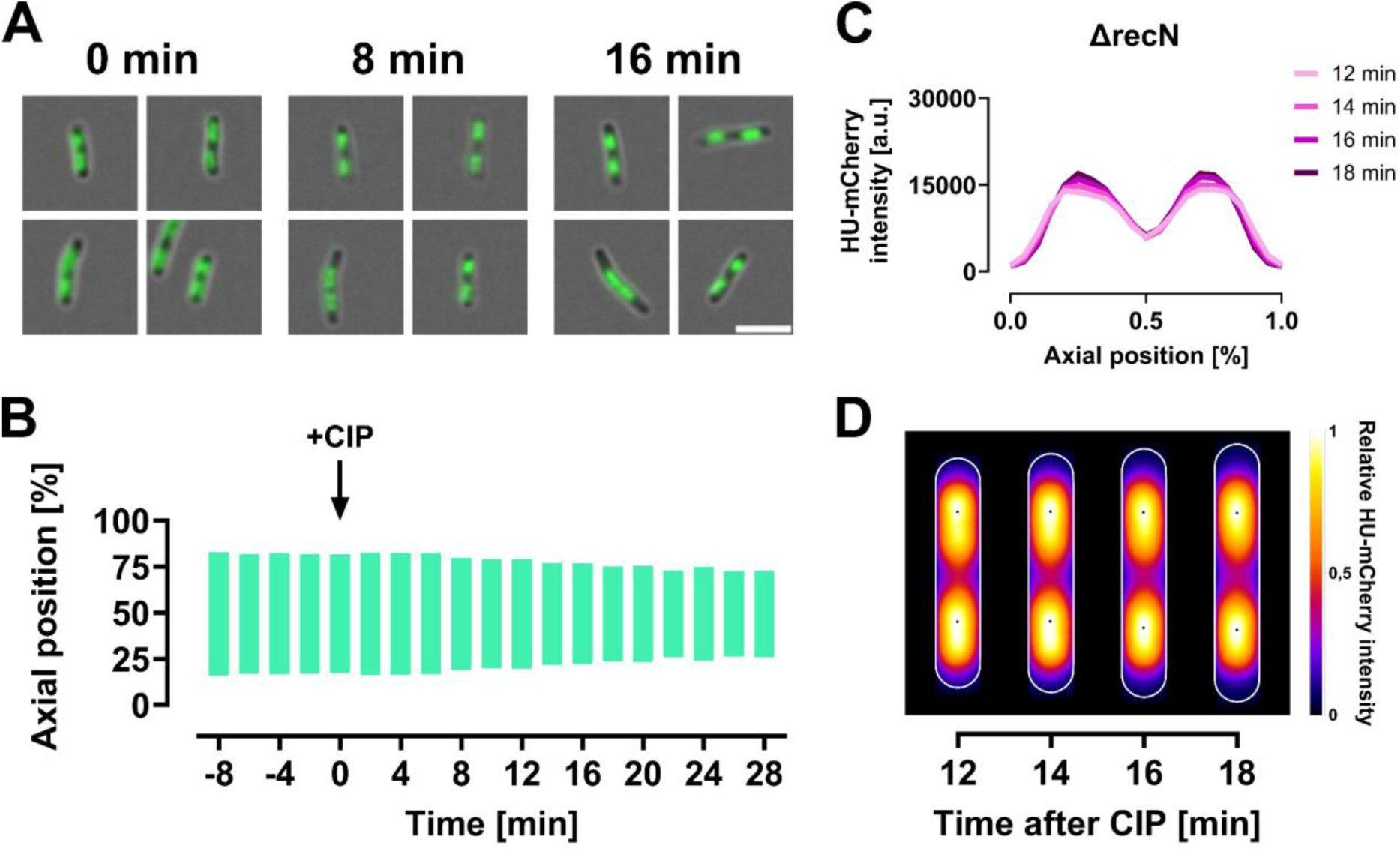
*ΔrecN* cells (MR16) exhibit limited nucleoid reorganization after CIP exposure. All cells were grown in LB at 37°C and imaged at 2-minute intervals using live-cell imaging. (**A** and **B**) Cells were immobilized in microfluidic channel slides and imaged using widefield microscopy. (**A**) Representative images of *ΔrecN* cells at 0, 8, and 16 minutes after CIP exposure, showing HU-mCherry fluorescence (green) to represent DNA distribution. Scale bar: 5 µm. (**B**) Analysis of DNA distribution along the cells’ long axis before and after CIP, quantified by measuring the distance between the outer bounds of symmetrical fluorescence peaks at 80% of maximum averaged intensity for each time point. Results are averaged from 17-61 cells from a single representative biological replicate (see Materials and methods for detailed explanation). (**C** and **D**) Cells immobilized on agar pads and imaged using spinning disk microscopy. Results are shown from images captured 12, 14, 16, and 18 minutes after CIP exposure, averaged from 40 tracked cells from a single representative biological replicate. (**C**) Average HU-mCherry fluorescence intensity along the cells’ long axis. (**D**) Kymograph heat map of relative HU-mCherry intensity distribution over time. A.u., arbitrary unit.

Analyses of axial distribution and heat maps of HU-mCherry fluorescence from 40 tracked cells confirmed the non-responsive phenotype of *ΔrecN* cells (Figure 2C and D). These findings led us to conclude that RecN is essential for the CIP-induced DNA supercompaction process.

### RecN forms dynamic foci that migrate with the nucleoid towards midcell during DNA supercompaction

Previous studies have found that fluorescently tagged RecN typically forms foci at the cell poles or between nucleoids at midcell under unchallenged conditions (20, 33). Following DNA damage by MMC, a majority of RecN foci were found to localize at the nucleoid (20, 33). Given our discovery that RecN is essential for DNA supercompaction, we aimed to investigate its localization and dynamics after CIP exposure. To achieve this, we employed two low-copy number plasmids to express GFP-tagged RecN: pTF271 with an arabinose-inducible promoter (denoted pARA), and pSG101 with the native SOS-inducible RecN promoter (denoted pSOS). The pARA plasmid allowed us to study GFP-RecN dynamics in the earliest stages of the SOS response, while the pSOS plasmid ensured a native, timely expression of GFP-RecN throughout the SOS response (26). We introduced these plasmids into the wild-type strain harboring chromosomally encoded HU-mCherry (KV21). Cells carrying the pARA plasmid (KV61) were exposed to arabinose (0.05%) for 60 minutes before CIP addition and live-cell spinning disk microscopy.

As expected, under unchallenged conditions, GFP-RecN predominantly formed static foci at cell poles and occasionally at the center of dividing cells when expressed from pARA (Supplementary Figure S1A and C). After CIP exposure, we observed a striking change: RecN foci at the poles migrated towards the nucleoids in a dynamic manner during DNA supercompaction (Figure 3A and Video 2). Analysis of fluorescence intensity from 328-480 cells revealed that GFP-RecN migration coincided with the onset of DNA supercompaction (Figure 3C). Upon completion of supercompaction, the mean GFP-RecN fluorescence was evenly distributed between poles and the midcell nucleoid (Figure 3C). Single-cell tracking showed that this wide distribution reflected the dynamic movement of RecN between poles and nucleoid occurring after DNA supercompaction was completed (Figure 3E).

**Figure 3.**
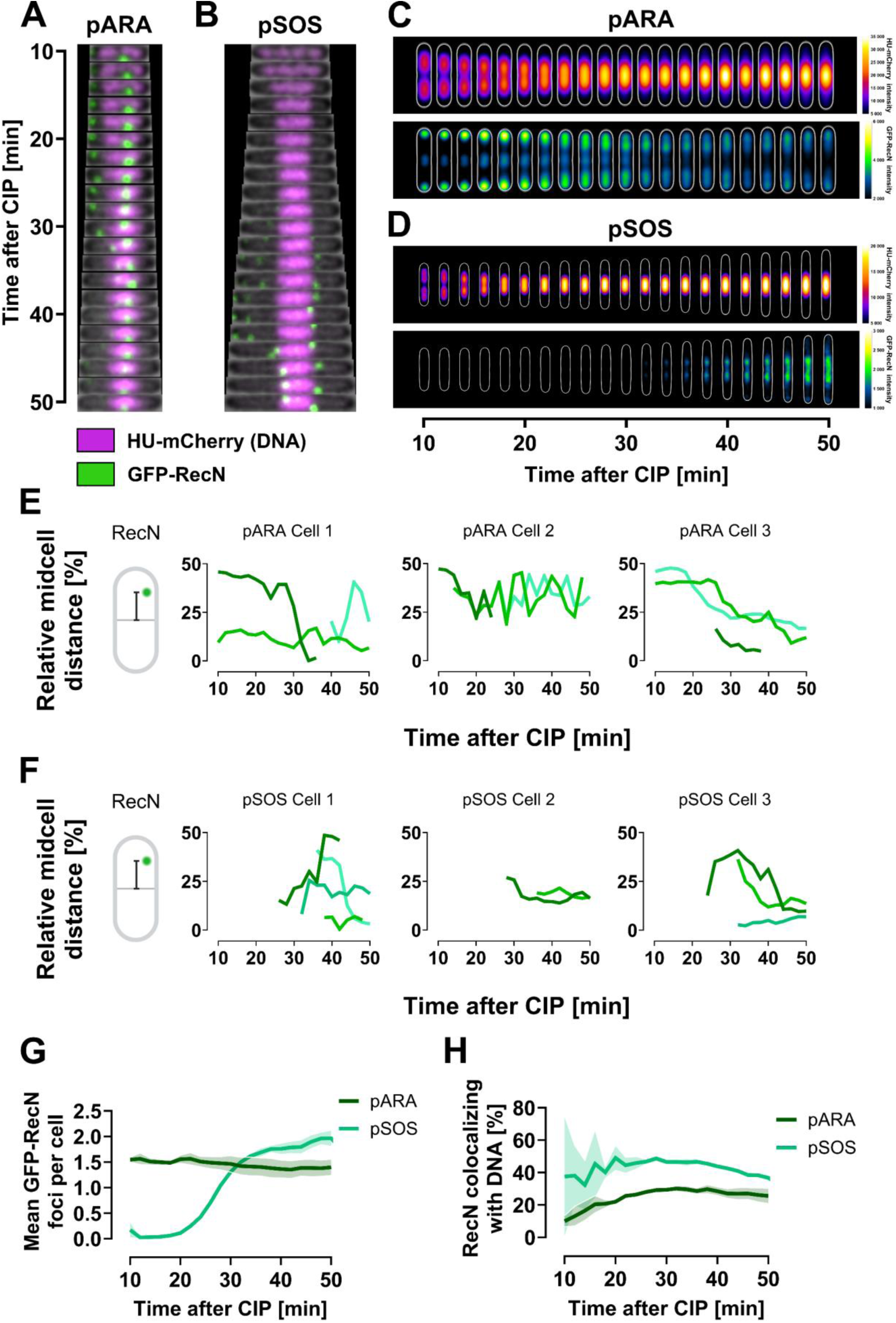
GFP-RecN foci exhibit dynamic movements in wild-type cells after CIP exposure. All cells were grown in LB at 37°C and imaged at 2-minute intervals using live-cell spinning disk microscopy starting 10 minutes post-CIP exposure. Results represent three biological replicates. (**A** and **B**) Kymographs of representative cells showing GFP-RecN dynamics in relation to DNA organization when GFP-RecN is expressed from (**A**) pARA (KV61) or (**B**) pSOS (KV60). GFP-RecN fluorescence is shown in green, while DNA is represented by HU-mCherry fluorescence in magenta. (**C** and **D**) Kymograph heat map of HU-mCherry intensity distribution (upper panels) and GFP-RecN intensity distribution (lower panels) inside cells over time, when GFP-RecN is expressed from (**C**) pARA (KV61) and (**D**) pSOS (KV60). Results at different time points are averaged from 1092-1612 cells for the pARA strain (KV61), and from 328-480 cells for the pSOS strain (KV60), in both cases from single representative biological replicates (see Materials and methods for detailed explanation). (**E** and **F**) GFP-RecN trajectories from representative cells showing the distance of GFP-RecN foci to midcell relative to cell length, when GFP-RecN is expressed from (**E**) pARA (KV61) and (**F**) pSOS (KV60). Each shade of green indicates a tracked focus. Midcell distance illustrations created in BioRender (https://BioRender.com/y04v494). (**G**) Percentage of GFP-RecN foci colocalizing with DNA versus time after CIP exposure for pARA and pSOS cells (KV61 and KV60). (**H**) Mean number of GFP-RecN foci per cell versus time after CIP exposure for pARA and pSOS cells (KV61 and KV60). Lines represent means from three biological replicates, and shaded regions indicate SEM.

**Video 2.**
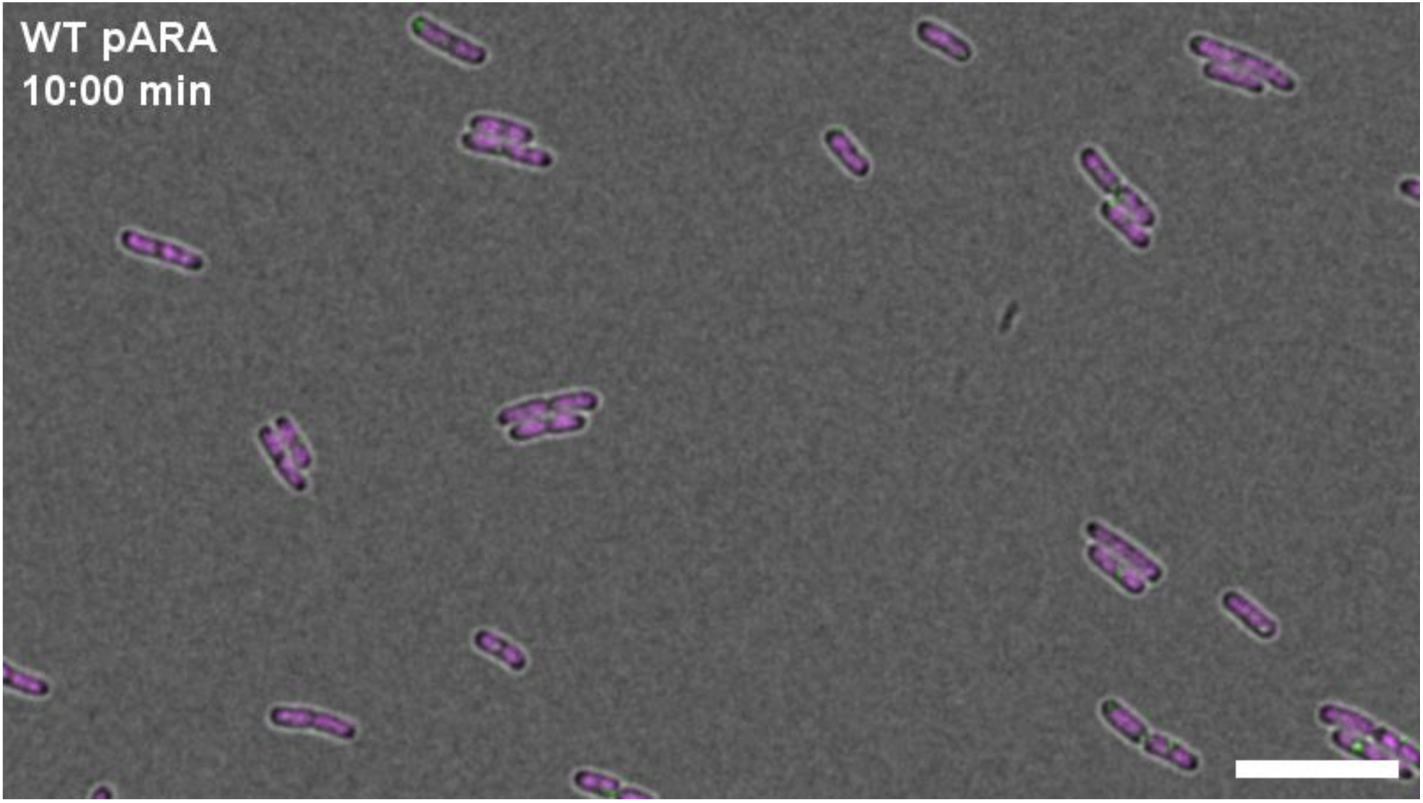
GFP-RecN dynamics in wild-type cells with pARA plasmid (KV61) after CIP exposure. GFP-RecN fluorescence is shown in green, while DNA is represented by HU-mCherry fluorescence in magenta. Cells were grown in LB at 37°C and imaged at 10-second intervals using live-cell spinning disk microscopy, starting 10 minutes after CIP exposure. Scale bar is 10 µm. [See Video 2 in Supplementary Data]

**Video 3.**
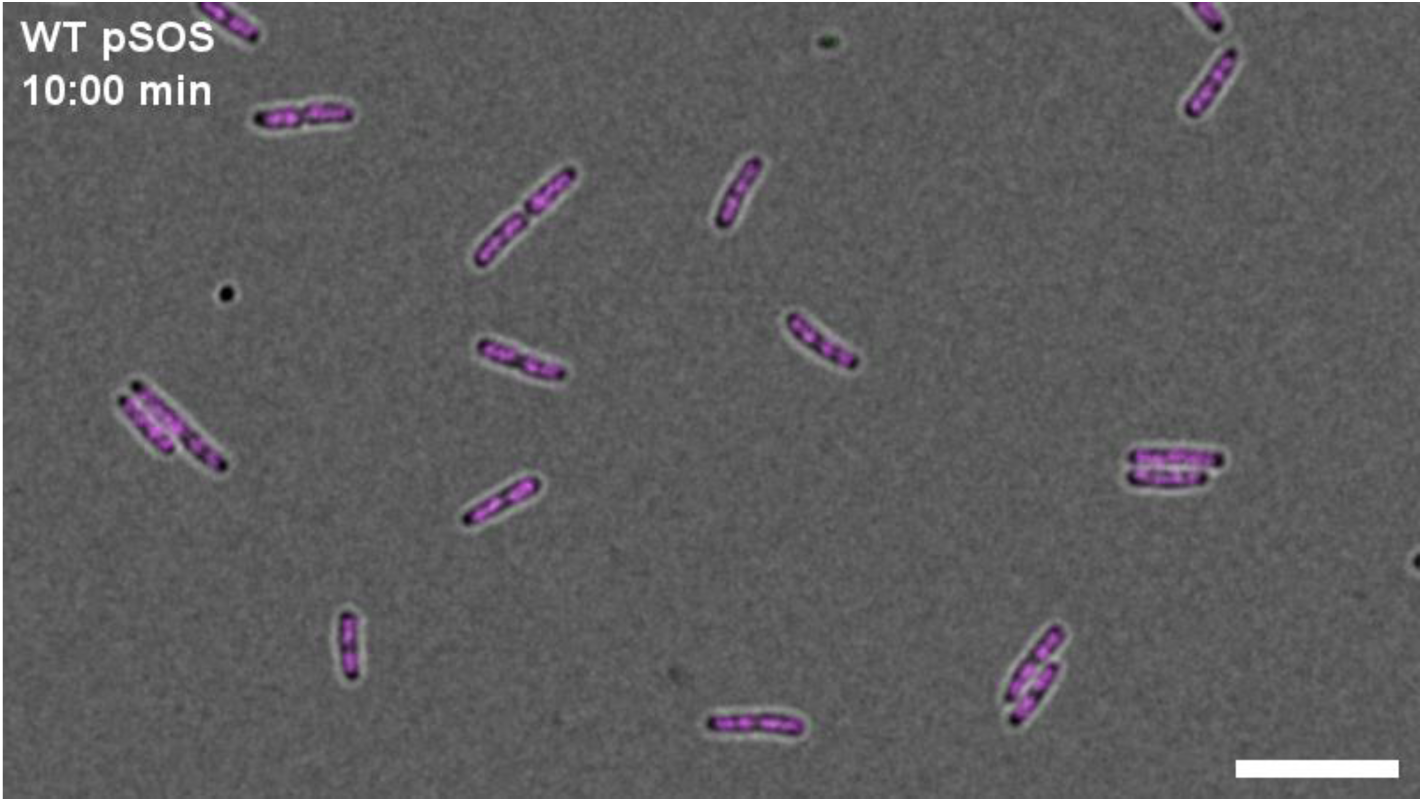
GFP-RecN dynamics in wild-type cells with pSOS plasmid (KV60) after CIP exposure. GFP-RecN fluorescence is shown in green, while DNA is represented by HU-mCherry fluorescence in magenta. Cells were grown in LB at 37°C and imaged at 10-seconds intervals using live-cell spinning disk microscopy, starting 10 minutes after CIP exposure. Scale bar is 10 µm. [See Video 3 in Supplementary Data]

In wild-type cells expressing GFP-RecN from the pSOS plasmid (KV60), almost no RecN foci were detectable at early time points after CIP exposure (Figure 3B and D). Foci first appeared around 30 minutes after CIP exposure, by which time DNA supercompaction was already complete. At this point, the pSOS cells reached the same mean number of GFP-RecN foci per cell as the pARA cells, before eventually surpassing it (Figure 3G). Furthermore, the pSOS cells exhibited dynamic GFP-RecN foci moving between poles and nucleoid once the foci appeared (Figure 3B and F, and Video 3), which aligned with observations from the pARA cells.

Comparing heatmaps from pARA and pSOS cells (Figure 3C and D), we found indications that the pSOS cells had a higher proportion of nucleoid-associated GFP-RecN foci than the pARA cells. Indeed, analysis of RecN and DNA colocalization revealed a consistently higher fraction of GFP-RecN foci localized at the nucleoid in the pSOS cells compared to the pARA cells (Figure 3H). Additionally, the pSOS cells exhibited faster DNA supercompaction than the pARA cells. Although DNA supercompaction started 12-14 minutes after CIP exposure in both strains, the pARA cells took about twice as long as the pSOS cells to complete supercompaction (Figure 3C and D, upper panels). These findings suggest that the premature expression or lack of temporal regulation of GFP-RecN in pARA cells influences DNA supercompaction dynamics and RecN localization.

We also introduced both plasmids into a *ΔrecN* background expressing HU-mCherry and found that the absence of endogenous *recN* slowed DNA supercompaction for both pSOS and pARA cells compared to in the wild-type background (Supplementary Figure S2A-D vs Figure 3). Furthermore, GFP-RecN foci from both plasmids in the *ΔrecN* background distributed more evenly across the cell after supercompaction than in the wild-type background (Supplementary Figure S2C and D, lower panels). The mean number of GFP-RecN foci per cell was considerably reduced for the pSOS cells (Supplementary Figure S2G), whereas overall colocalization of GFP-RecN with DNA increased for the pARA cells but decreased for the pSOS cells relative to the wild-type background (Supplementary Figure S2H). These findings may imply that GFP-RecN is less efficient than native RecN in DNA supercompaction and that native RecN proteins may be required within the GFP-RecN complexes for foci formation and correct function.

### GFP-RecN is functional in DNA supercompaction

The physiological relevance of GFP-RecN has been debated, with suggestions that foci at poles are aggregates not involved in biological activities (26). Indeed, we observe anomalies likely caused by GFP-RecN expression, such as slower DNA supercompaction, especially in pARA cells (Figure 3C) and exacerbated in a *ΔrecN* background (Supplementary Figure S2). However, GFP-RecN still drives DNA supercompaction in cells lacking native RecN (Supplementary Figure S2), though not as efficiently as when native RecN is present. The delay in DNA supercompaction is presumably linked to the cells’ overall RecN levels, or to the ratio between highly efficient native RecN and less efficient GFP-RecN.

To survey native RecN and GFP-RecN levels in our strains, we performed Western blotting. We compared protein levels in pARA– and pSOS-containing strains with their wild-type or *ΔrecN* background strains, both before and after 20 minutes of CIP exposure (Supplementary Figure S3A and B). The pARA strains were incubated with arabinose (0.05%) for 60 minutes before CIP addition. Our results showed that overall RecN levels in pSOS and pARA strains were only modestly increased by approximately 2-3 and 4 times, respectively, compared to the wild-type background without plasmid, after subtracting the signal from the *ΔrecN* negative control (Supplementary Figure S3C). The *ΔrecN* strains carrying pSOS exhibited RecN levels comparable to that of the wild-type.

We next examined if GFP-RecN presence and increased overall RecN levels affect survival by performing spot assays using strains carrying either the pARA or pSOS plasmids and comparing them with their background strains. We found that these plasmids did not negatively affect survival rates in either wild type or *ΔrecN* backgrounds after 20 minutes of CIP exposure (Supplementary Figure S4). Additionally, GFP-RecN expression in *ΔrecN* backgrounds restored survival to wild-type levels. These results are consistent with previous findings in MMC-treated cells (20, 33).

Despite elevated overall RecN levels during DNA supercompaction in cells with pARA or pSOS plasmids, GFP-RecN still drives supercompaction, restores survival to wild-type levels, and exhibit dynamic movement between poles and supercompacted DNA. This indicates that GFP-RecN is functionally active in RecN-dependent processes, while potentially performing better when native RecN is present.

### RecA is essential for DNA supercompaction and RecN dynamics

Having established that GFP-RecN can drive DNA supercompaction, we next investigated the role of RecA in the dynamics of RecN. Previous studies showed that fluorescently tagged RecA, similar to GFP-RecN, typically localizes at cell poles under unchallenged conditions and migrates to the nucleoid after DNA damage (33). In addition, RecN and RecA colocalize in nucleoid gaps after MMC treatment in *E. coli* (33), and RecA dynamics depend on RecN in *Caulobacter crescentus* and *Bacillus subtilis* (34, 55). Based on these insights, we hypothesized that RecN dynamics after CIP exposure depend on RecA. To test this hypothesis, we performed live-cell imaging of GFP-RecN in a *ΔrecA* background harboring HU-mCherry.

Strikingly, in the *ΔrecA* background, GFP-RecN expression was insufficient to facilitate DNA supercompaction after CIP exposure (Figure 4A and Video 4). It was instead observed that many cells merged their DNA at midcell while it remained distributed across the cell length (Figure 4C). Furthermore, GFP-RecN foci remained at cell poles or in nucleoid gaps after CIP exposure (Figure 4A and C, and Video 4). Single-cell analyses revealed diminished movement of GFP-RecN across the cell length in the *ΔrecA* background compared to the wild type (Figure 4E vs Figure 3E). As expected, *ΔrecA* cells harboring the pSOS plasmid did not produce any GFP-RecN foci because of the absence of SOS induction in this background (Figure 4B and D). The findings above are consistent with observations in a *ΔrecN ΔrecA* background (Supplementary Figure S5).

**Figure 4.**
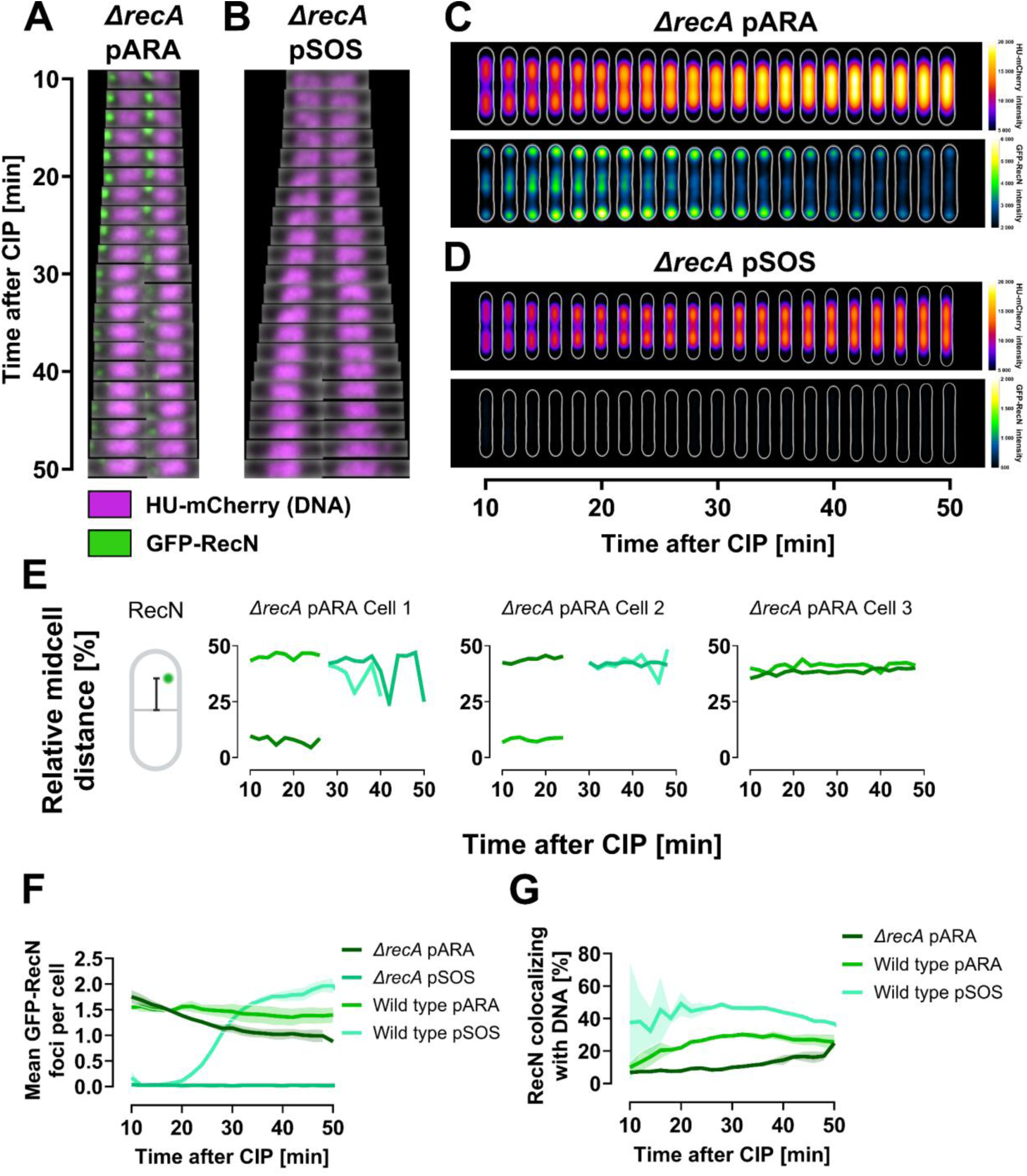
GFP-RecN foci dynamics after CIP exposure are diminished in *ΔrecA* cells. All cells were grown in LB at 37°C and imaged at 2-minute intervals using live-cell spinning disk microscopy starting 10 minutes post-CIP exposure. Results represent three biological replicates. (**A** and **B**) Kymographs of representative *ΔrecA* cells showing GFP-RecN dynamics and DNA organization when GFP-RecN is expressed from (**A**) pARA (KV67) or (**B**) pSOS (KV66). GFP-RecN fluorescence is shown in green, while DNA is represented by HU-mCherry fluorescence in magenta. (**C** and **D**) Kymograph heat map of HU-mCherry intensity distribution (upper panels) and GFP-RecN intensity distribution (lower panels) inside cells over time, when GFP-RecN is expressed from (**C**) pARA (KV67) and (**D**) pSOS (KV66). Results at different time points are averaged from 579-899 cells for the *ΔrecA* pARA strain (KV67), and from 513-894 cells for the *ΔrecA* pSOS strain (KV66), in both cases from single representative biological replicates (see Materials and methods for detailed explanation). (**E**) GFP-RecN trajectories from representative *ΔrecA* pARA cells (KV67) showing the distance of GFP-RecN foci to midcell relative to cell length. Each shade of green indicates a tracked focus. Midcell distance illustration created in BioRender (https://BioRender.com/y04v494). (**F**) Percentage of GFP-RecN foci colocalizing with DNA versus time after CIP exposure for *ΔrecA* pARA and pSOS cells (KV67 and KV66). (**G**) Mean number of GFP-RecN foci per cell versus time after CIP exposure for *ΔrecA* pARA and pSOS cells (KV67 and KV66). Lines represent means from three biological replicates, and shaded regions indicate SEM.

**Video 4.**
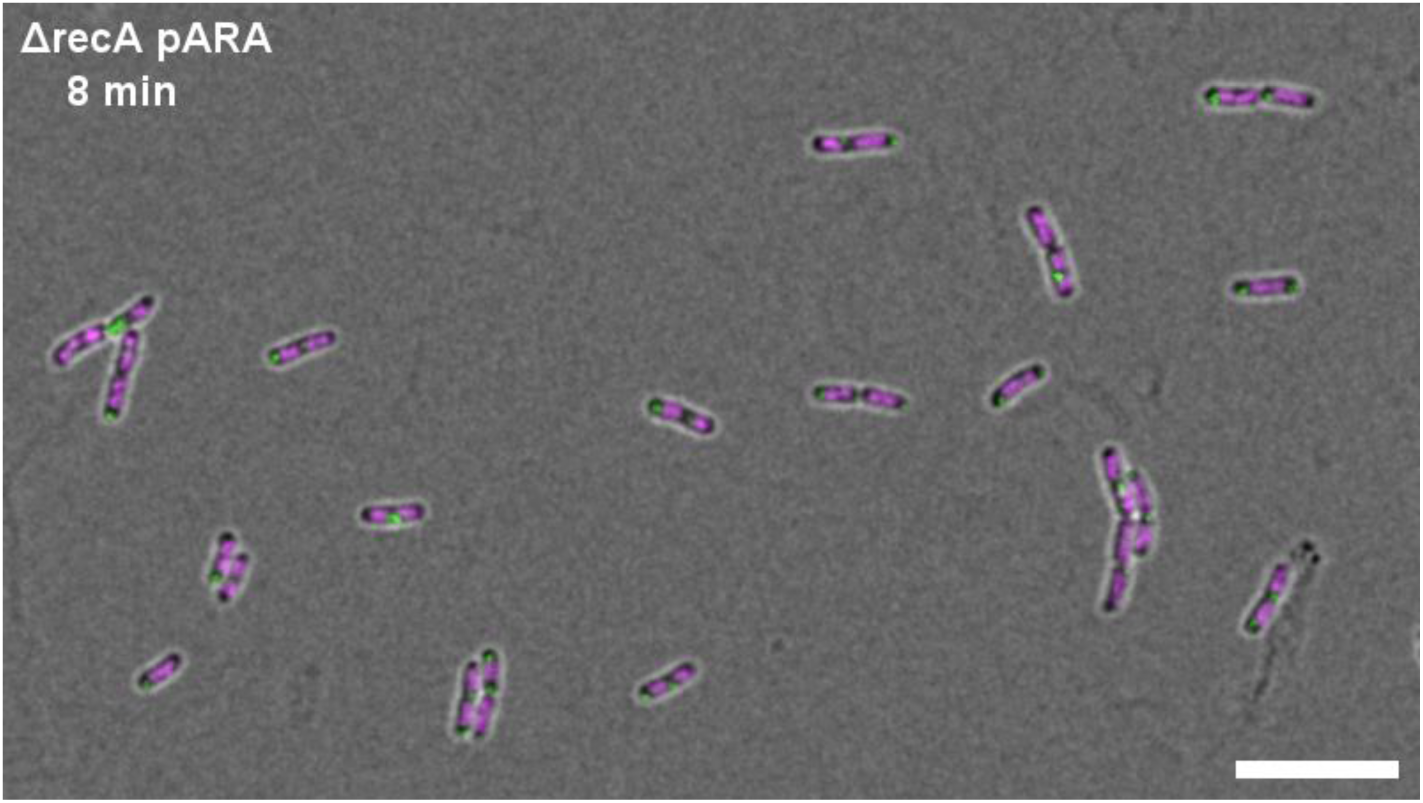
GFP-RecN dynamics in ΔrecA cells with pARA plasmid (KV67) after CIP exposure. GFP-RecN fluorescence is shown in green, while DNA is represented by HU-mCherry fluorescence in magenta. Cells were grown in LB at 37°C and imaged at 2-minute intervals using live-cell spinning disk microscopy, starting 8 minutes after CIP exposure. Scale bar is 10 µm. [See Video 4 in Supplementary Data]

The mean number of arabinose-induced GFP-RecN foci remained largely unaffected by the absence of RecA during DNA supercompaction (Figure 4F). However, the proportion of these foci colocalizing with DNA considerably decreased in the absence of RecA compared to wild-type or *ΔrecN* backgrounds (Figure 4G), aligning with a previous report (20). This reduced colocalization with DNA cannot be attributed to a deficiency in the formation of GFP-RecN foci.

In summary, the absence of RecA abolishes DNA supercompaction, even when GFP-RecN is artificially expressed from a plasmid. Additionally, it disrupts RecN localization and diminishes RecN dynamics. These findings indicate that RecN cannot function properly without either RecA, induction of the SOS response, or both.

### Functional RecA is necessary for timely DNA supercompaction

Our investigation thus far established that RecN cannot function properly in DNA supercompaction without RecA. However, it remains unclear whether the necessity of RecA stems only from its role as an SOS response inducer or from a direct interplay with RecN, as others have proposed (19, 20, 32–34, 55, 56). To differentiate between these roles, we examined strains with temperature-regulated SOS induction, possessing either native RecA (*recA^+^*) or non-functional RecA (*recA1*). The strains contained two mutations in the lexA gene — *lexA41 (lexA3)* — which make LexA repressor cleavage, and thus SOS induction, depend solely on the temperature-sensitive activity of Lon protease rather than RecA (44).

At low temperature (30°C), which corresponds to low Lon activity and presumably low expression of RecN, both *recA^+^* and *recA1* cells (KP84 and DM936) failed to exhibit DNA supercompaction within the expected 20 minutes after CIP exposure (Figure 5A 30°C, and B-C). The *recA1* cells struggled to exhibit DNA supercompaction even within 60 minutes, while the *recA^+^* cells achieved supercompaction within 40 minutes (Figure 5A 30°C). Thus, although supercompaction was delayed in both strains, the delay was more prominent in cells without functional RecA.

**Figure 5.**
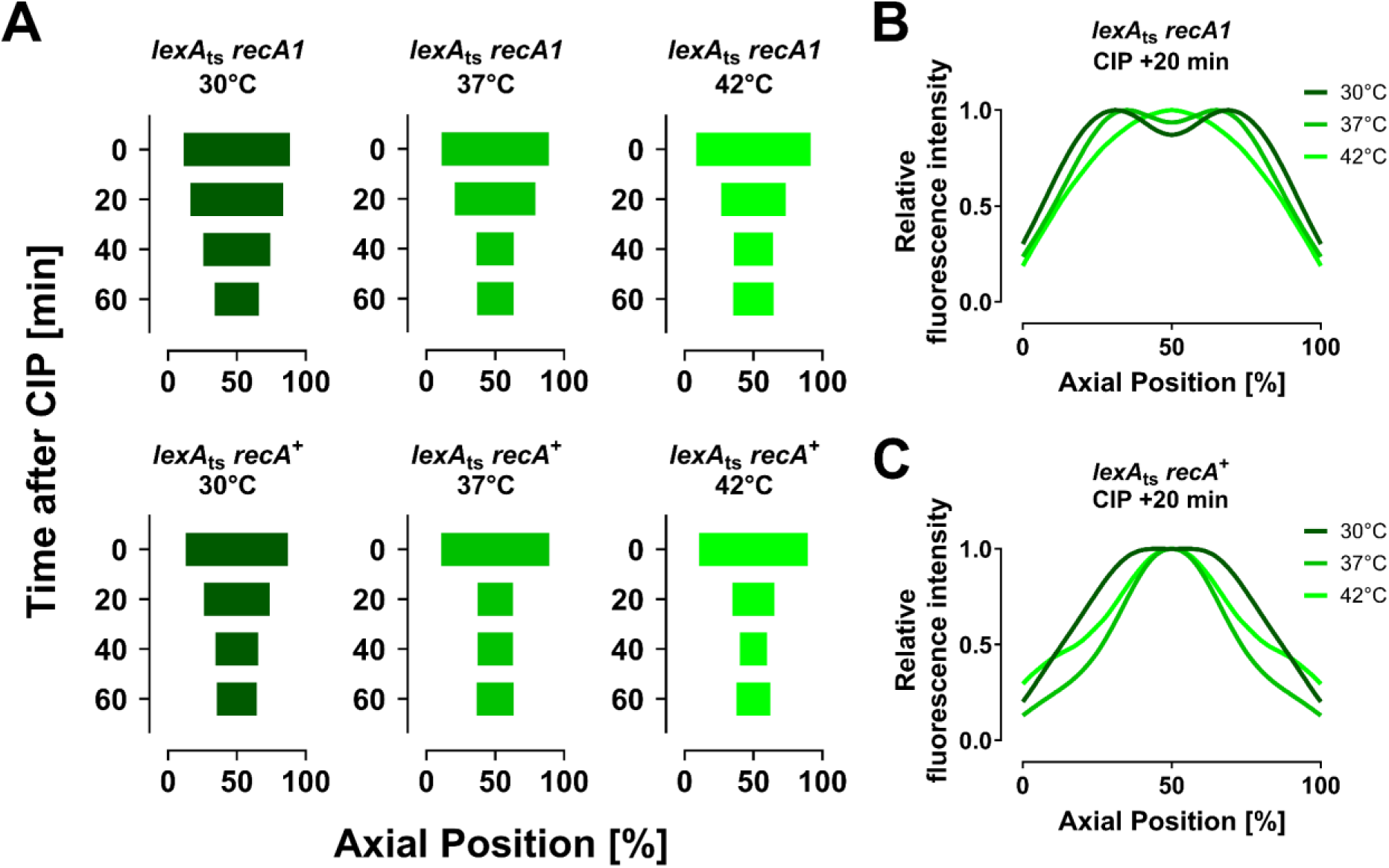
Effect of RecA functionality on DNA distribution following CIP exposure in cells with temperature-regulated SOS induction (*lexA*_ts_). Comparison between strain (KP84) with native RecA (*recA*^+^) and strain (DM936) with non-functional RecA (*recA1*). Cells were grown in LB and exposed to CIP at the indicated temperatures before fixation. Fixed cells were stained with Hoechst 33258 for DNA visualization. Results for each time point are averaged from 550-8546 cells from a single representative biological replicate (see Materials and methods for detailed explanation). (**A**) Analysis of DNA distribution along the cells’ long axis before and 20, 40, and 60 minutes after CIP for (upper panel) cells with non-functional RecA (DM936) and (lower panel) cells with native RecA (KP84). DNA distribution is quantified by measuring the distance between the outer bounds of fluorescence peaks at 80% of maximum averaged intensity for each time point. (**B** and **C**) Normalized average Hoechst fluorescence intensity along the cells’ long axis 20 minutes after CIP exposure for (**B**) cells with non-functional RecA (DM936) and (**C**) cells with native RecA (KP84).

At elevated temperatures (37°C and 42°C) and increased Lon activity, the *recA^+^* cells exhibited DNA supercompaction within the expected 20 minutes after CIP (Figure 5A bottom row, and C). In contrast, the *recA1* cells still failed to complete supercompaction within this wild-type time frame (Figure 5A top row, and B). The clear difference in supercompaction timing between the two strains indicates that the functional RecA is more essential for DNA supercompaction following CIP exposure than the induction of the SOS response itself.

In summary, while SOS response induction contributes to DNA supercompaction and is necessary for RecN expression, functional RecA is in itself essential for a timely response to severe DNA damage from CIP. RecA’s essential role may possibly stem from a direct interplay with RecN, as suggested in previous studies (19, 30, 32).

### RecN and RecA interact and show frequent transient colocalization at nucleoid-associated positions during DNA supercompaction

Building on our observations of RecA’s essential role in DNA supercompaction, we aimed to elucidate the interplay between RecN and RecA. For this purpose, we performed live-cell spinning disk microscopy on cells expressing fluorescently tagged versions of both proteins. RecA-mCherry was expressed in tandem with the endogenous *recA* gene, while GFP-RecN was expressed from either the pARA or pSOS plasmid (KV79 and KV77). We confirmed that these strains exhibited DNA supercompaction after CIP exposure by imaging fixed cells stained with Hoechst 33258 (Supplementary Figure S6).

RecA-mCherry foci displayed rapid kinetics, often appearing and disintegrating swiftly (Figure 6A and B, and Video 5 and 6), which complicated tracking of individual foci to characterize their dynamics. These rapid kinetics are emphasized by the wide average distribution of RecA-mCherry across cells at all time points (Figure 6C and D, upper panels), and the noticeable variation in the mean number of foci per cell (Supplementary Figure S7A). Around 40 minutes after CIP exposure, single RecA-mCherry foci tended to disappear, while overall RecA-mCherry fluorescence increased and spread more evenly across the cells (Figure 6C and D, upper panels). Consequently, we centered our analysis on the cellular localization of RecA-mCherry foci after CIP exposure and their potential colocalization with GFP-RecN foci.

**Figure 6.**
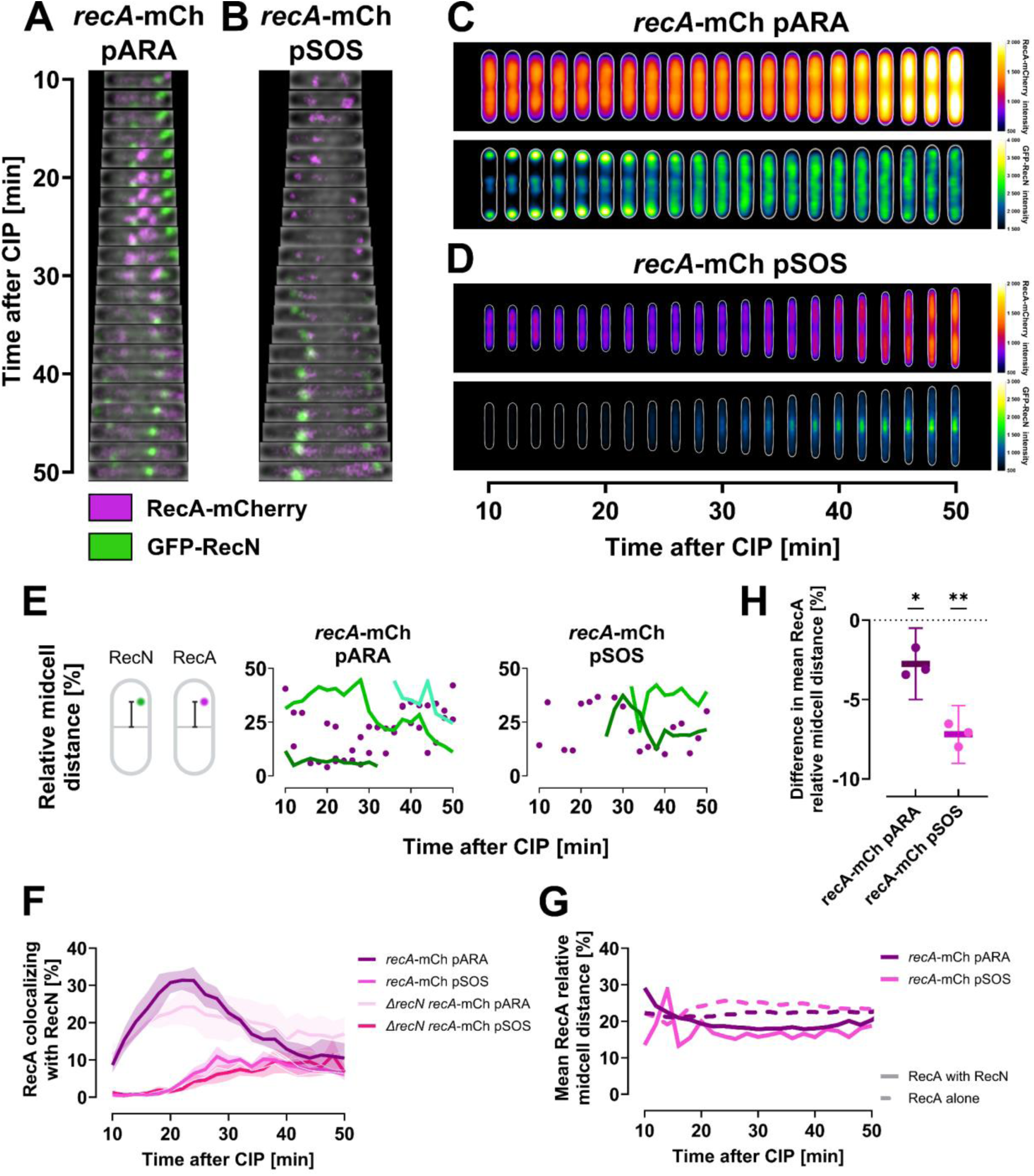
RecA-mCherry foci frequently and transiently colocalize with GFP-RecN foci after CIP exposure, with colocalizing RecA-mCherry foci positioned closer to midcell. All cells were grown in LB at 37°C, expressing RecA-mCherry chromosomally alongside native RecA. Cells were imaged at 2-minute intervals starting 10 minutes post-CIP exposure using live-cell spinning disk microscopy. Results represent three biological replicates. (**A**, **B**) Kymographs of representative wild-type cells showing RecA-mCherry (magenta) and GFP-RecN (green) dynamics when GFP-RecN is expressed from either (**A**) pARA (KV79) or (**B**) pSOS (KV77). (**C** and **D**) Kymograph heat map of RecA-mCherry intensity distribution (upper panels) and GFP-RecN intensity distribution (lower panels) inside cells over time, when GFP-RecN is expressed from (**C**) pARA (KV79) or (**D**) pSOS (KV77). Results at different time points are averaged from 480-792 cells for the pARA strain (KV79), and from 231-398 cells for the pSOS strain (KV77), in both cases from single representative biological replicates (see Materials and methods for detailed explanation). (**E**) GFP-RecN trajectories and RecA-mCherry localization for representative cells expressing GFP-RecN from pARA (KV79, left) and pSOS (KV77, right). Plots display the distance of foci to midcell relative to cell length, with GFP-RecN trajectories in shades of green and RecA-mCherry foci as magenta dots. Midcell distance illustrations created in BioRender (https://BioRender.com/y04v494). (**F**) Percentage of RecA-mCherry foci colocalizing with GFP-RecN foci versus time after CIP exposure for wild-type and *ΔrecN* cells with pARA or pSOS plasmids. Lines represent means from three biological replicates, and shaded regions indicate SEM. (**G**) Mean distance of colocalizing (solid line) and non-colocalizing (dashed line) RecA-mCherry foci to midcell relative to cell length. Means are from three biological replicates. (**H**) Difference in relative midcell distance for colocalizing versus non-colocalizing RecA-mCherry foci. Negative values indicate that RecA-mCherry foci colocalizing with GFP-RecN are closer to midcell than non-colocalizing RecA-mCherry. Lines show means, dots represent replicates’ mean values, and error bars indicate 95% CIs; * *p* ≤ 0.05; ** *p* ≤ 0.01.

We observed that RecA-mCherry frequently and transiently colocalized with GFP-RecN (Figure 6E and F, Videos 5 and 6, and Supplementary Figure S8). In the pARA-carrying cells, we noted a remarkable spike in RecA-mCherry colocalizing with GFP-RecN, which coincided with the period of DNA supercompaction (Figure 6F). The pSOS cells, with barely any detectable GFP-RecN foci during DNA supercompaction, did not exhibit a similar colocalization spike (Supplementary Figure S7B). Nevertheless, after 40 minutes of CIP exposure, the pSOS cells achieved colocalization levels similar to the wild-type pARA cells (Figure 6F).

Additionally, RecA-mCherry foci colocalizing with GFP-RecN were located significantly closer to midcell, and thus closer to the supercompacted DNA, than non-colocalizing RecA-mCherry foci in both pARA and pSOS cells (Figure 6G and H). Colocalizing foci were 2.8% (95% CI 0.5-5.0; *p*=0.03) and 7.2% (95% CI 5.4-9.0; *p*=0.003) closer to midcell relative to cell length for wild-type pARA and pSOS cells, respectively. This proximity to midcell became particularly prominent after more than 20 minutes of CIP exposure, corresponding to the completion of DNA supercompaction (Figure 6G and Supplementary Figure S6). Lack of endogenous *recN* reduced but did not eliminate the midcell proximity differences in both pARA and pSOS cells (Supplementary Figure S7C and D). Using a bacterial two hybrid assay, we also demonstrated a strong direct interaction between RecN and RecA (Supplementary Figure S9 and Table S2), consistent with earlier studies (19, 30, 32). In summary, these findings indicate that RecN and RecA interact and function cooperatively in proximity of DNA during supercompaction.

**Video 5.**
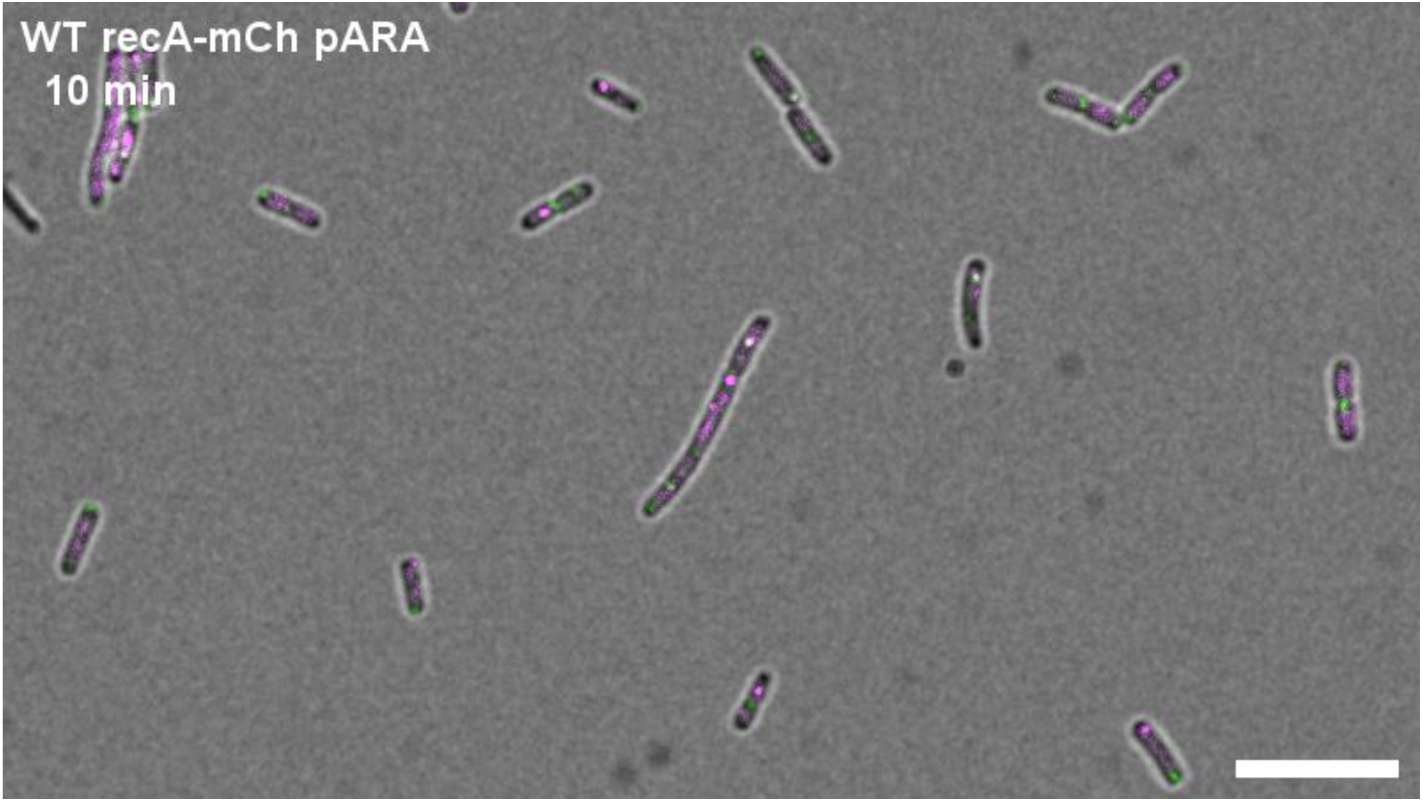
RecA-mCherry and GFP-RecN dynamics in wild-type cells with chromosomally expressed recA-mCherry and pARA plasmid (KV79) after CIP exposure. RecA-mCherry fluorescence is shown in magenta, while GFP-RecN fluorescence is shown in green. Cells were grown in LB at 37°C and imaged at 2-minute intervals using live-cell spinning disk microscopy, starting 10 minutes after CIP exposure. Scale bar is 10 µm. [See Video 5 in Supplementary Data]

**Video 6.**
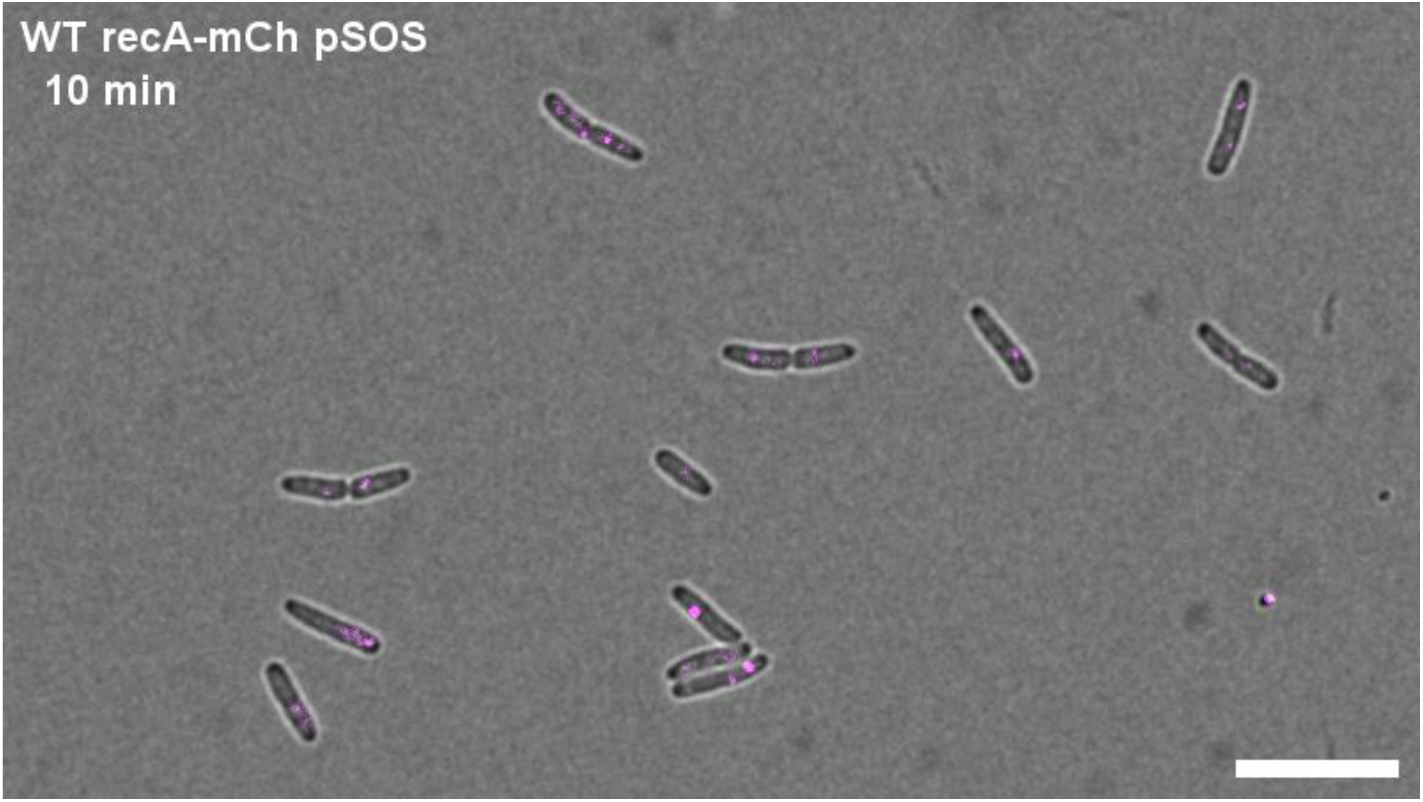
RecA-mCherry and GFP-RecN dynamics in wild-type cells with chromosomally expressed recA-mCherry and pSOS plasmid (KV77) after CIP exposure. RecA-mCherry fluorescence is shown in magenta, while GFP-RecN fluorescence is shown in green. Cells were grown in LB at 37°C and imaged at 2-minute intervals using live-cell spinning disk microscopy, starting 10 minutes after CIP exposure. Scale bar is 10 µm. [See Video 6 in Supplementary Data]

### Absence of RecN sensitizes cells to ciprofloxacin

RecN and RecA are both involved in DSB repair (28), although the specific role of RecN is still under discussion. Building on our findings that RecN and RecA interact and possibly cooperate during DNA supercompaction, we aimed to assess the importance of RecN for survival following CIP-induced DNA damage at our experimental dose (10 µg/mL). We found that *ΔrecN* cells (JW5416) are extremely sensitive to short-term CIP exposure at this dose, with survival rates plummeting to the detection limit within just 1 minute (Figure 7). This increased sensitivity aligns with an essential role of RecN in both DNA supercompaction and DNA repair, emphasizing its critical function in the cellular response to CIP-induced DNA damage.

**Figure 7.**
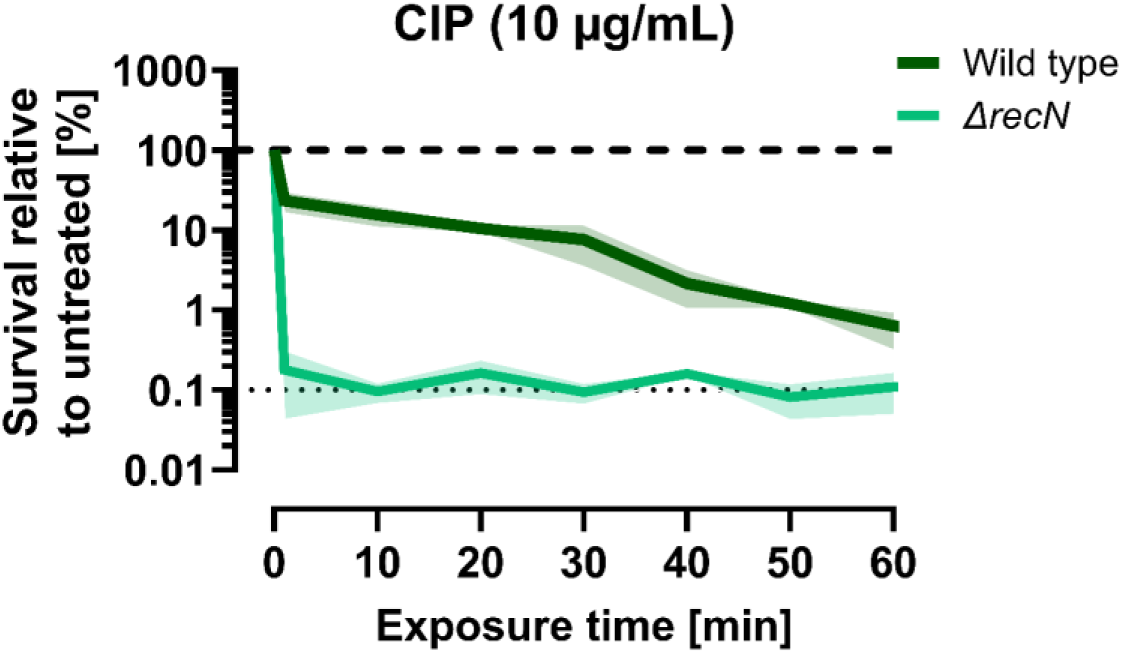
*ΔrecN* cells are overly sensitive to CIP. Time-dependent survival assay for wild-type (BW25113) and *ΔrecN* (JW5416) cells after exposure to 10 µg/mL CIP. Exposure time refers to the duration of CIP exposure before washing and spotting the cells. Survival was measured as colony forming units CFU/mL and relative survival was calculated by comparing with unchallenged parallels. Thick dashed line indicates survival of unchallenged parallel; thin dotted line indicates the assay’s detection limit. Green lines represent means of 3 biological replicates; shaded regions indicate standard deviation.

## DISCUSSION

In this study, we show that treatment of *E. coli* cells with the fluoroquinolone antibiotic CIP leads to a major reorganization of DNA into a dense structure at midcell — a process we term DNA supercompaction. Live-cell fluorescence microscopy reveals that this process occurs in a stepwise and highly organized manner, in which nucleoid lobes initially merge at quarter positions after which all DNA fuses and compacts at midcell (Figure 1). Given the uniformity of this process, we propose that DNA supercompaction is part of an active cellular response to severe DNA damage.

### RecN is the main actor in DNA supercompaction

We identify RecN as essential for DNA supercompaction in response to severe DNA damage caused by CIP. Cells lacking RecN fail to exhibit DNA supercompaction (Figure 2), consistent with earlier findings that RecN is required for DNA compaction following exposure to other DNA-damaging agents such as MMC (21), bleomycin (21), gamma-irradiation (53), and UV-irradiation (18). RecN is rapidly induced following DNA damage (13) and reaches high levels within 10 minutes of CIP exposure (24). However, it undergoes rapid degradation by ClpXP protease, with a half-life of around 10 minutes (26). This swift yet transient presence of RecN at elevated levels indicates a vital role in the immediate DNA damage response.

To shed more light on the activity of RecN in the DNA supercompaction process, we tracked the localization of functionally active GFP-tagged RecN. We discovered that RecN foci migrated with the nucleoids towards midcell during DNA supercompaction and subsequently moved dynamically between poles and midcell (Figure 3). RecN also frequently colocalized with the DNA (Figure 3H), indicating activity on the DNA. Combined with our finding that cells lacking RecN cannot perform DNA supercompaction (Figure 2), we suggest that RecN has an active role in reorganizing DNA after CIP exposure.

In vitro studies have shown that RecN can bind and bridge DNA molecules. Keyamura and Hishida demonstrated that purified *E. coli* RecN can bind both ssDNA and dsDNA, with a preference for ssDNA (30). They proposed that RecN is recruited to ssDNA at DSBs, where it can topologically entrap a second dsDNA in an ATP-dependent manner (30). Similar properties have been observed in vitro for *Deinococcus radiodurans* RecN, which can bridge dsDNA molecules through ATP-dependent activity (23, 31). While the relevance of these in vitro findings for in vivo functions remains to be fully elucidated, our results, supported by other studies (19, 33), show that RecN requires the presence of DSBs to perform DNA compaction (Supplementary Figure S1). This indicates that RecN’s activity in DNA compaction may be linked to the immediate DNA damage response.

### RecN activity in DNA supercompaction depends on RecA

RecA plays a crucial role in the repair of DSBs through its involvement in homology search and strand invasion (10, 57). RecN is also established as an actor within this pathway (27, 28), although its exact role is still not fully understood. In our study, we found that RecN depends on RecA to effectively facilitate DNA supercompaction (Figure 4). This dependency on RecA was not solely connected to its role in inducing the SOS response and thereby the expression of RecN or other repair proteins (Figure 5). Furthermore, we observed frequent colocalization of RecN and RecA at nucleoid-associated positions following CIP exposure (Figure 6) and found evidence supporting a direct interaction (Supplementary Figure S9 and Table S2).

Recent studies on the interplay between RecN and RecA following DNA damage have predominantly centered on the hypothesis that RecN ensures close proximity between damaged and intact homologous DNA, thereby facilitating RecA’s homology search and strand invasion (19, 20, 32–34, 55, 56). Efficient homology search likely requires both short-distance sampling through 1D sliding and hopping, and long-distance sampling through 3D intersegmental transfer on the DNA, depending on the location of the break(s) relative to intact homologous DNA (57, 58). Vickridge et al. showed that RecN facilitates interaction between newly replicated strands of homologous DNA when DSBs arise at the replication fork, in a manner dependent on RecA (19). In this case, the “zippering” activity of RecN could enable a rapid and effective homology search by RecA in the immediate vicinity of the DSB. In cases where DSBs are generated on already replicated and segregated DNA, RecA filaments appear to extend from the DSB throughout the cell, likely in the search for homologous DNA, as demonstrated in both *E. coli* and *C. crescentus* (34, 52, 54). It has been indicated that the highly dynamic behavior of RecA filaments depends on the ATPase activity of RecN (34), and that this activity of RecN is also necessary for successful RecA-mediated repair (20, 33).

In vitro studies with purified *D. radiodurans* RecN support the in vivo studies and highlight the co-dependency between RecN and RecA. It was found that RecA’s function in DSB repair is dependent on the ATPase activity of RecN, and that ssDNA-bound RecA filaments heavily stimulate the ATPase activity of RecN (32). It was suggested that this increased level of ATPase activity could power the movement of the RecN-RecA-ssDNA complex in the search for homologous DNA at distant sites (32). Additionally, the confined, dense nucleoid may in itself increase efficiency of homology search between distant sites of the DNA (57, 59).

Although we cannot conclude mechanistic details concerning the RecA-RecN interaction from our study, it is evident that RecN and RecA exhibit a co-dependency not only in DSB repair but also in DNA supercompaction. The increased RecN-RecA colocalization we observed during DNA supercompaction (Figure 6F) could possibly imply that DNA supercompaction is associated with a spike in RecN ATPase activity (32). Furthermore, the close proximity of colocalized RecA-RecN foci to midcell (Figure 6G and H) may indicate cooperation at or near the compacted DNA.

### DNA supercompaction is likely a consequence of severe DNA damage

The morphology and dynamics of bacterial nucleoids has been investigated using different genotoxic agents under varying conditions, which might lead to discrepancies between reported results. In our study, the cells likely harbored up to 24 active replication forks, depending on cell cycle stage (60), and were exposed to a clinically relevant dose of CIP (10 µg/mL) about 500 times the MIC. This dose likely leads to severe DNA damage through generation of numerous DSBs at the replication forks and other locations of the DNA (3). We found that DNA supercompaction occurred within 8-20 minutes of exposure (Figure 1). This type of extensive DNA compaction has also been observed after exposure to nalidixic acid, another quinolone antibiotic (17). In contrast, mild UV exposure was found to lead to a more transient and irregularly timed DNA compaction (18, 61). Furthermore, upon induction of only a single DSB, no DNA compaction was observed (52). Instead, only the DSB-containing region and the region with homologous DNA were transported to midcell for repair and then redistributed to the original positions (52). These findings may indicate that the DNA compaction phenotype and its kinetics depend on the severity of DNA damage and the number or type of lesions needing repair.

To further explore this possibility, we analyzed the nucleoid dynamics and survival of wild-type (BW25113) and *ΔrecN* (JW5416) cells after exposure to a milder CIP dose (MIC, 20 ng/mL). Consistent with our hypothesis, we observed a much slower DNA compaction at this dose, as the cell population did not exhibit notable DNA compaction until after approximately 60 minutes (Supplementary Figure S10A). The midcell nucleoid structure observed after 60 minutes also appeared less dense than when using the higher CIP dose (Supplementary Figure S10A vs Figure 1C). The *ΔrecN* cells failed to exhibit DNA supercompaction, as expected (Supplementary Figure S10B). Correspondingly, the cells also required significantly longer exposure time to CIP at this dose before the increased sensitivity of *ΔrecN* cell became prominent (Supplementary Figure S10C).

Interestingly, Meddows et al. found that RecN is more important for cell survival after the induction of three well-separated DSBs than after a single DSB (27). In contrast, cells lacking RecA showed equal sensitivity to both one and three DSBs (27), indicating a more prominent role of RecN when multiple DSBs need repair. As previously discussed, RecA’s stimulation of RecN’s ATPase activity is important for entrapment of secondary dsDNA in vitro and “zippering” of newly replicated DNA in vivo. Evidence further indicates an intricate interplay between RecN and RecA in homology search and repair. In our situation with numerous DSBs, a presumably large number of RecN proteins binding and bridging DNA in an attempt to repair the severe damage, could lead to the observed DNA supercompaction phenotype (62).

### Functional and structural parallels between RecN and Eukaryotic SMC proteins in compaction and repair

As an SMC-like protein, RecN belongs to a group of conserved proteins that are considered the primary drivers of DNA organization and compaction (63, 64). Eukaryotic cells use multiple SMC complexes for tight chromosome organization, including the SMC5/6 complex (65). Recent reports on SMC5/6 reveal roles and activities that are strikingly similar to those discussed for RecN. The SMC5/6 complex facilitates DSB repair, stabilizes arrested replication forks, and has a high affinity for ssDNA-dsDNA junctions and other recombination intermediates (66–68). The complex can slide along the DNA and dimerize at junctions relevant for homologous recombination, and is then thought to compact damaged DNA regions by loop extrusion in an ATP-dependent manner (see (65) for review). Given the functional and structural resemblances between RecN and SMC5/6, it may be that they operate through similar mechanisms.

In conclusion, our study reveals that RecN, in concert with RecA, plays an essential role in the highly organized DNA supercompaction following severe DNA damage caused by CIP in *E. coli*. We propose that DNA compaction is part of a universal response to DSBs, with its degree dependent on the severity of the damage. Although a role of RecN in aiding RecA homology search could explain this effect, further investigation is needed. Given the rising prevalence of antibiotic resistance, particularly to fluoroquinolones, insights into bacterial survival responses to DNA-damaging antibiotics are crucial for developing preventive measures and novel treatments to combat resistance.

## DATA AVAILABILITY

The data underlying this article will be shared on reasonable request to the corresponding author. All scripts and templates used for processing and analysis of images in this study are available in the Zenodo repository at https://doi.org/10.5281/zenodo.14063054.

## SUPPLEMENTARY DATA

Supplementary Data are available online at bioRxiv.

## AUTHOR CONTRIBUTIONS

Conceptualization: K.V., J.B., K.S., E.H.; Methodology: K.V., S.B.R., I.M.R., J.B., K.S., E.H; Investigation: K.V., S.B.R, I.M.R.; Formal analysis: K.V., I.M.R.; Visualization: K.V.; Software: K.V.; Funding acquisition: K.V., M.B., J.B., K.S., E.H.; Supervision: J.B., K.S., E.H.; Writing — original draft: K.V., E.H.; Writing — review & editing: all authors.

## ACKNOWLEDGEMENTS

We thank David Leach (University of Edinburgh) for the pDL5196 plasmid containing the *recA-mCherry* construct, Takashi Hishida (Gakushuin University) for the GFP-RecN plasmids (pARA and pSOS), and Steven Sandler (University of Massachusetts at Amherst) for the SS6282 strain carrying HU-mCherry. We also thank Adrien Ducret (Université de Lyon) for assistance with MicrobeJ analysis and code development, and Thierry Oms (Université Libre de Bruxelles) for assistance with kymograph plots. Microscopy was performed at the Advanced Light Microscopy core facility, Gaustad node at Oslo University Hospital, Gaustad. The graphical abstract was created in BioRender (https://BioRender.com/s57i772). We acknowledge the assistance of ChatGPT by OpenAI in providing suggestions on phrasing and suggesting code for scripts. The human authors performed all editing.

## TABLE AND FIGURES LEGENDS

**Supplementary Table S1.** Plasmids used for the bacterial two hybrid assay in this study.

**Supplementary Figure S1.** DNA distribution and GFP-RecN dynamics in wild-type and *ΔrecN* cells under unchallenged conditions.

**Supplementary Figure S2.** GFP-RecN is functionally active and drives DNA supercompaction in *ΔrecN* cells, although supercompaction is slower without native RecN.

**Supplementary Figure S3.** Western blots with anti-RecN antibody showing the effect of GFP-RecN expression from pARA and pSOS on overall levels of RecN.

**Supplementary Figure S4**. Spot assay of strains with pSOS and pARA plasmids to evaluate the effect of GFP-RecN expression on survival after CIP.

**Supplementary Figure S5.** GFP-RecN dynamics in *ΔrecN ΔrecA* cells after CIP exposure.

**Supplementary Figure S6.** Analysis of DNA distribution inside all strains with chromosomally expressed RecA-mCherry to evaluate whether this expression alone or in combination with GFP-RecN affects DNA supercompaction.

**Supplementary Figure S7.** Foci count over time for RecA-mCherry and GFP-RecN foci in wild-type and *ΔrecN* strains, and analysis of RecA-mCherry midcell distance in *ΔrecN* background.

**Supplementary Figure S8.** Examples of GFP-RecN trajectories and RecA-mCherry localization for representative cells from strains with chromosomally expressed RecA-mCherry.

**Supplementary Figure S9.** Relative β-galactosidase activity results from bacterial two hybrid assay of RecN and RecA interaction.

**Supplementary Table S2.** Relative β-galactosidase activity results from bacterial two hybrid assay of RecN and RecA interaction.

**Supplementary Figure S10.** *ΔrecN* cells are overly sensitive to CIP also at the MIC dose and fail to compact DNA like wild-type cells within 60 minutes after exposure to MIC doses of CIP.

