## Supplementary Data for "RecN and RecA orchestrate an ordered DNA supercompaction response following ciprofloxacin exposure in *Escherichia coli*"

**Table S1.** Plasmids used for the bacterial two hybrid assay in this study. $Amp^R$ = Ampicillin resistance; $Kan^R$ = Kanamycin resistance; MCS, Multiple cloning site.

| Plasmid | Description | Source |
|---|---|---|
| pKT25/pKNT25 | pSU40 derivative encoding T25-fragment (first 224 amino acids of *cyaA*) downstream (pKT25) or upstream (pKNT25) of the MCS [$Kan^R$] | Euromedex (cat. no. EUK001) |
| pUT18/pUT18C | pUC19 derivative encoding T18-fragment (amino acids 225-399 of *cyaA*) downstream (pUT18) or upstream (pUT18C) of the MCS [$Amp^R$] | Euromedex (cat. no. EUK001) |
| pKT25-zip | pKT25 harboring T25 genetically fused to the leucine zipper of GCN4 [$Kan^R$] | Euromedex (cat. no. EUK001) |
| pUT18C-zip | pUT18C harboring T18 genetically fused to the leucine zipper of GCN4 [$Amp^R$] | Euromedex (cat. no. EUK001) |
| pKT25-*recA* | pKT25 with *recA* inserted in frame with T25 in MCS (BamHI and EcoRI restriction sites) [$Kan^R$] | GenScript |
| pKNT25-*recA* | pKNT25 with *recA* inserted in frame with T25 in MCS (BamHI and EcoRI restriction sites) [$Kan^R$] | GenScript |
| pKT25-*recN* | pKT25 with *recN* inserted in frame with T25 in MCS (BamHI and EcoRI restriction sites) [$Kan^R$] | GenScript |

| | | |
|---|---|---|
| pKNT25-*recN* | pKNT25 with *recN* inserted in frame with T25 in MCS (BamHI and EcoRI restriction sites) [$Kan^R$] | GenScript |
| pUT18-*recA* | pUT18 with *recA* inserted in frame with T18 in MCS (BamHI and EcoRI restriction sites) [$Amp^R$] | GenScript |
| pUT18C-*recA* | pUT18C with *recA* inserted in frame with T18 in MCS (BamHI and EcoRI restriction sites) [$Amp^R$] | GenScript |
| pUT18-*recN* | pUT18 with *recN* inserted in frame with T18 in MCS (BamHI and EcoRI restriction sites) [$Amp^R$] | GenScript |
| pUT18C-*recN* | pUT18C with *recA* inserted in frame with T18 in MCS (BamHI and EcoRI restriction sites) [$Amp^R$] | GenScript |

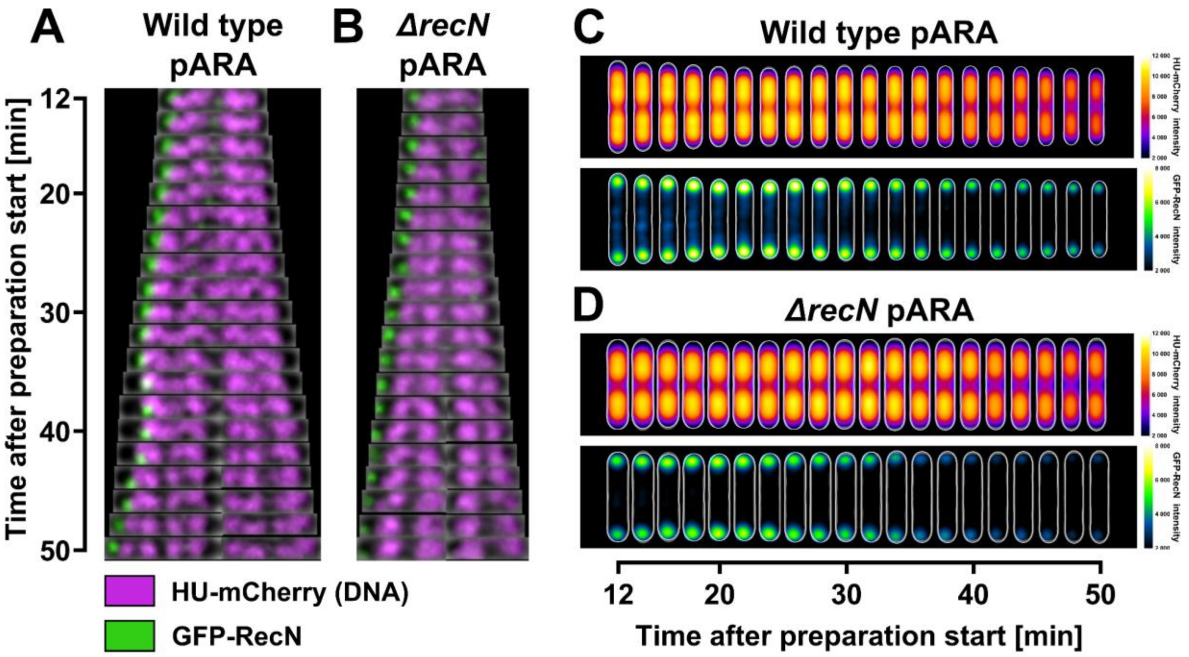


**Figure S1.** DNA distribution and GFP-RecN dynamics in wild-type and *ΔrecN* cells under unchallenged conditions. All cells were grown in LB at 37°C and imaged at 2-minute intervals using live-cell spinning disk microscopy starting 12 minutes after preparation on agar pads. (**A** and **B**) Kymographs of representative cells showing GFP-RecN dynamics in relation to DNA organization under unchallenged conditions in (**A**) wild-type pARA (KV61) cells and (**B**) *ΔrecN* pARA (KV63) cells. GFP-RecN fluorescence is shown in green, while DNA is represented by HU-mCherry fluorescence in magenta. (**C** and **D**) Kymograph heat map of HU-mCherry intensity distribution (upper panels) and GFP-RecN intensity distribution (lower panels) inside cells over time, under unchallenged conditions in (**C**) wild-type pARA (KV61) cells and (**D**) *ΔrecN* pARA (KV63) cells. Results at different time points are averaged from 446-1255 cells for the wild-type pARA strain (KV61), and from 638-1183 cells for the *ΔrecN* pARA strain (KV63), in both cases from single representative biological replicates (see Materials and methods for detailed explanation).

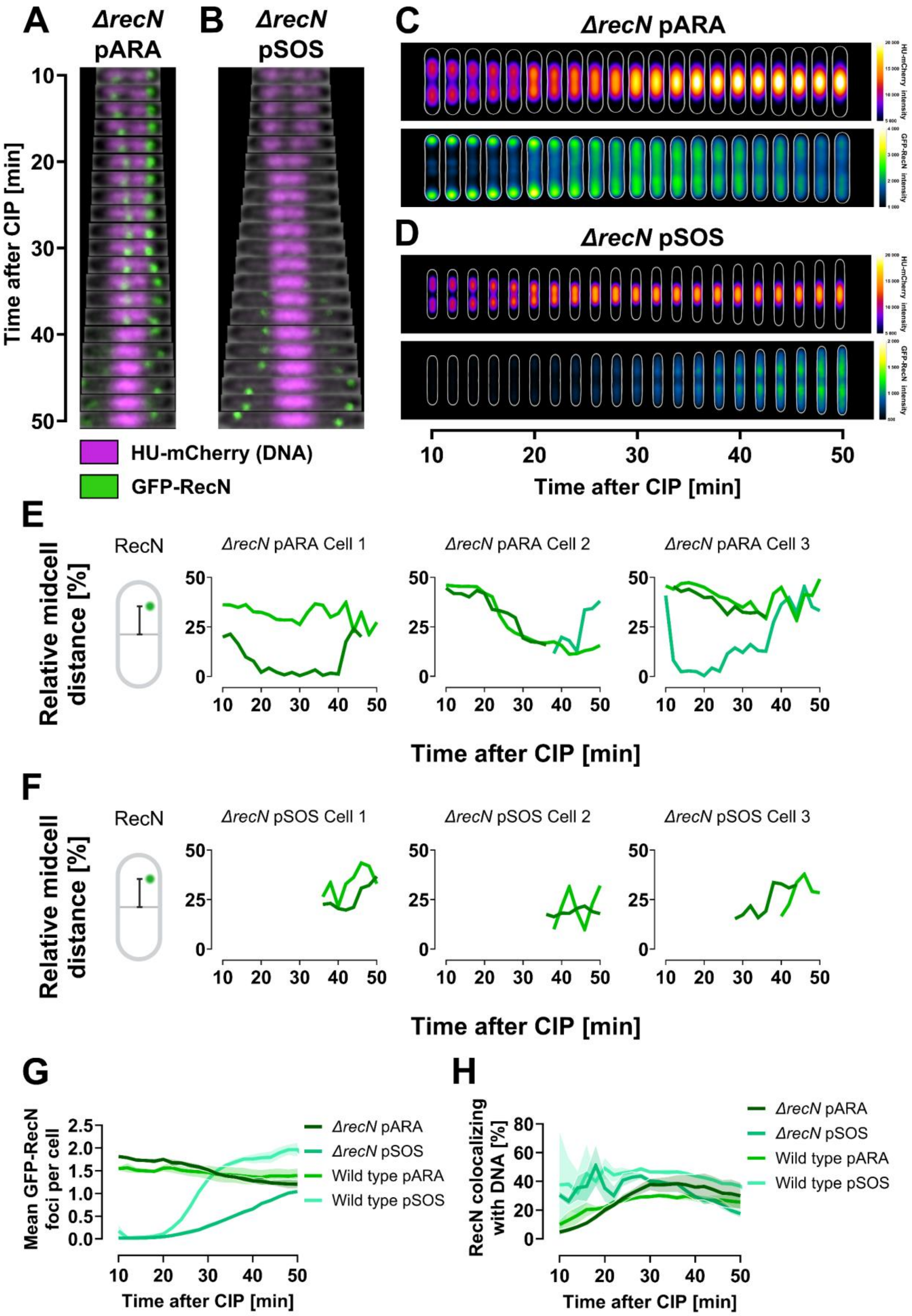


*Figure S2 caption on next page*

**Figure S2.** GFP-RecN is functionally active and drives DNA supercompaction in *ΔrecN* cells, although supercompaction is slower without native RecN. All cells were grown in LB at 37°C and imaged at 2-minute intervals using live-cell spinning disk microscopy starting 10 minutes post-CIP exposure. Results represent three biological replicates. (**A** and **B**) Kymographs of representative *ΔrecN* cells showing GFP-RecN dynamics in relation to DNA organization when GFP-RecN is expressed from (**A**) pARA (KV63) or (**B**) pSOS (KV62). GFP-RecN fluorescence is shown in green, while DNA is represented by HU-mCherry fluorescence in magenta. (**C** and **D**) Kymograph heat map of HU-mCherry intensity distribution (upper panels) and GFP-RecN intensity distribution (lower panels) inside cells over time, when GFP-RecN is expressed from (**C**) pARA (KV63) and (**D**) pSOS (KV62). Results at different time points are averaged from 711-1082 cells for the *ΔrecN* pARA strain (KV63), and from 445-775 cells for the *ΔrecN* pSOS strain (KV62), in both cases from single representative biological replicates (see Materials and methods for detailed explanation). (**E** and **F**) GFP-RecN trajectories from representative *ΔrecN* cells showing the distance of GFP-RecN foci to midcell relative to cell length, when GFP-RecN is expressed from (**E**) pARA (KV63) and (**F**) pSOS (KV62). Each shade of green indicates a tracked focus. Midcell distance illustrations created in BioRender (https://BioRender.com/y04v494). (**G**) Percentage of GFP-RecN foci colocalizing with DNA versus time after CIP exposure for pARA and pSOS cells (KV63 and KV62). (**H**) Mean number of GFP-RecN foci per cell versus time after CIP exposure for *ΔrecN* pARA and pSOS cells (KV63 and KV62). Lines represent means from three biological replicates, and shaded regions indicate SEM.

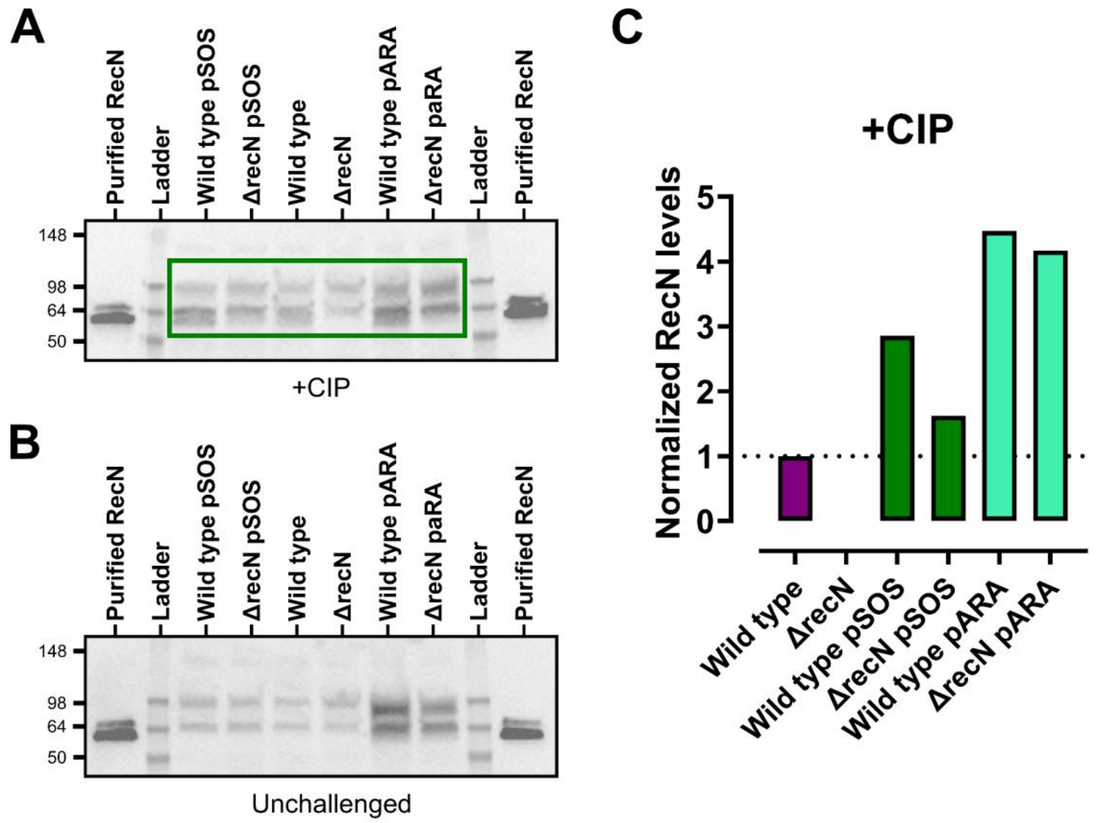


**Figure S3.** Western blots with anti-RecN antibody showing the effect of GFP-RecN expression from pARA and pSOS on overall levels of RecN. (**A** and **B**) Wild-type and *ΔrecN* cells without GFP-RecN plasmids (middle lanes) were used as controls for RecN levels; purified RecN (outer lanes) were used as controls for the antibody; ladders (second outmost lanes) are Invitrogen SeeBlue Plus2 Pre-stained Protein Standard and protein sizes noted on left edge of western blots. Strains were grown to exponential phase in LB at 37°C, and strains with pARA plasmids were incubated with arabinose (0.05%) for 60 minutes. All strains were then prepared for western blots either (**A**) after 20 minutes of CIP exposure, or (**B**) under unchallenged conditions. (**C**) Overall RecN protein levels normalized to the wild-type strain (dotted line) for each lane inside the green box in (A). The quantification simultaneously compares the overall levels of both native RecN (61 kDa) and GFP-RecN (98 kDa) between the different strains.

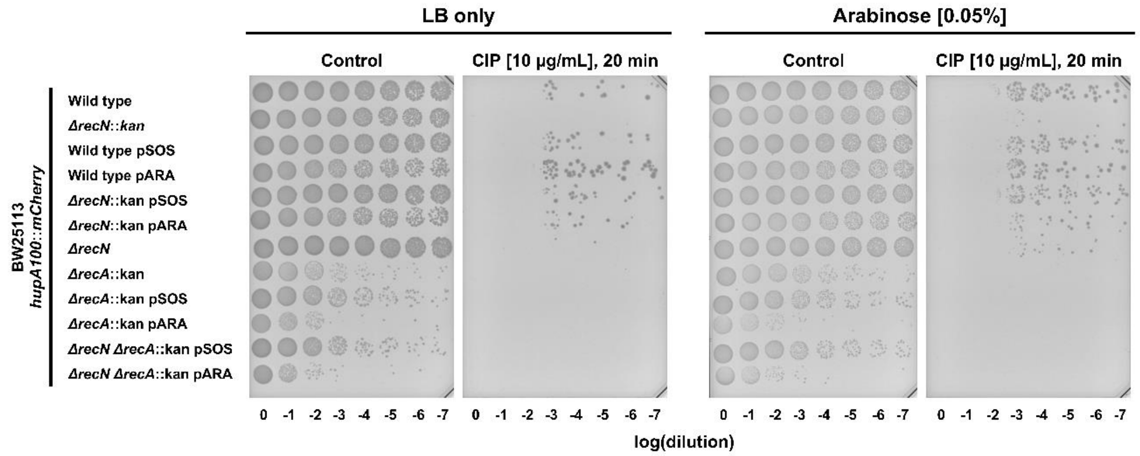


**Figure S4.** Spot assay of strains with pSOS and pARA plasmids to evaluate the effect of GFP-RecN expression on survival after CIP. These backgrounds include wild type with chromosomally expressed HU-mCherry (KV21), as well as *ΔrecN*, *ΔrecA*, and *ΔrecN ΔrecA* variants. All strains were grown to exponential phase in LB at 37°C. Strains were then either exposed to CIP (10 µg/mL) for 20 minutes or left unchallenged (control). Cultures were 10-fold serial diluted, as indicated along the x-axis. Finally, the diluted cultures were plated out on LB-agar plates with or without arabinose (0.05%). CIP concentration is too high for survival in dilutions below 1:1000.

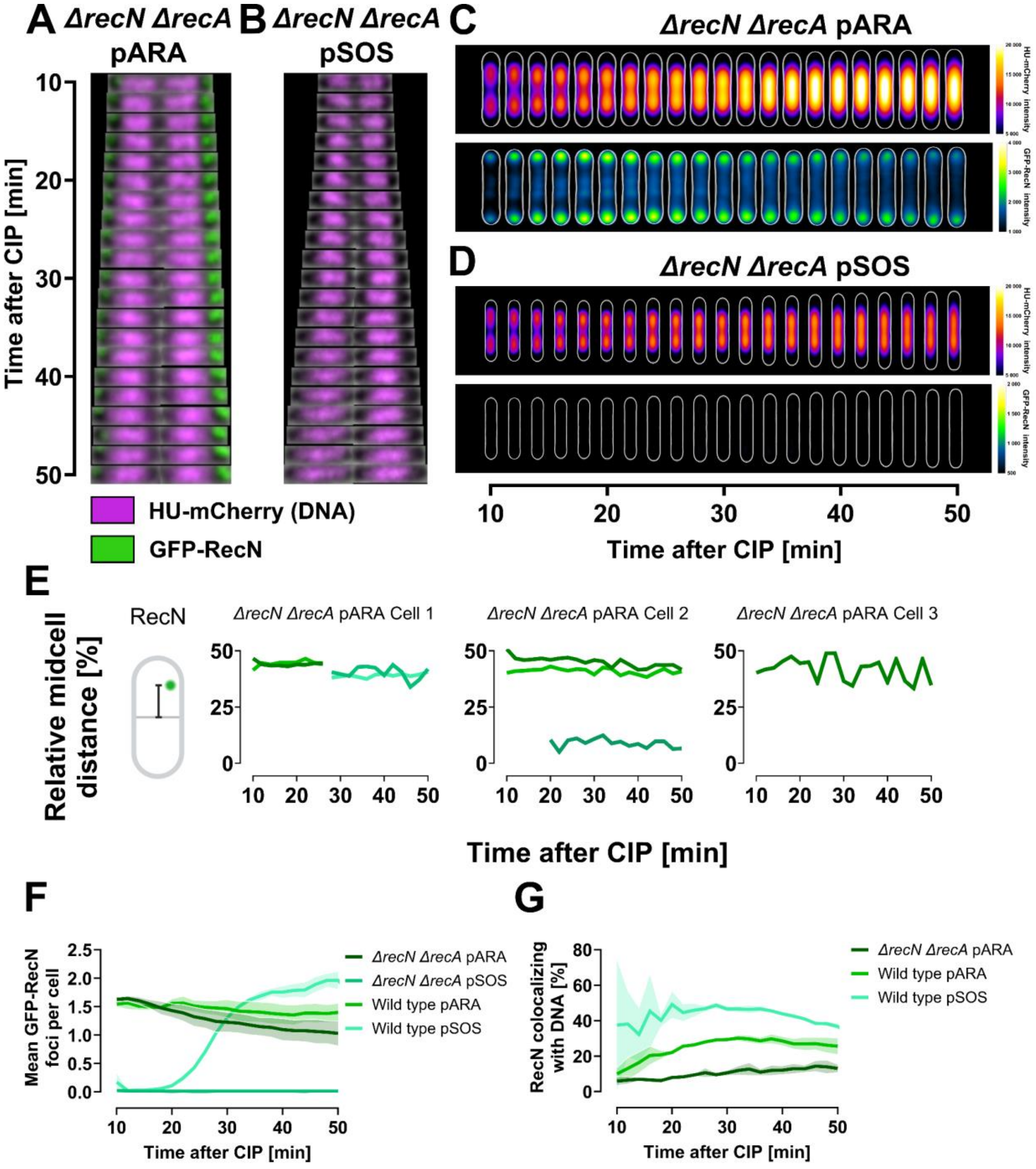


*Figure S5 caption on next page*

**Figure S5.** GFP-RecN dynamics in *ΔrecN ΔrecA* cells after CIP exposure. All cells were grown in LB at 37°C and imaged at 2-minute intervals using live-cell spinning disk microscopy starting 10 minutes post-CIP exposure. Results represent three biological replicates. (**A** and **B**) Kymographs of representative *ΔrecN ΔrecA* cells showing GFP-RecN dynamics and DNA organization when GFP-RecN is expressed from (**A**) pARA (KV71) or (**B**) pSOS (KV70). GFP-RecN fluorescence is shown in green, while DNA is represented by HU-mCherry fluorescence in magenta. (**C** and **D**) Kymograph heat map of HU-mCherry intensity distribution (upper panels) and GFP-RecN intensity distribution (lower panels) inside cells over time, when GFP-RecN is expressed from (**C**) pARA (KV71) and (**D**) pSOS (KV70). Results at different time points are averaged from 545-953 cells for the *ΔrecN ΔrecA* pARA strain (KV71), and from 335-539 cells for the *ΔrecN ΔrecA* pSOS strain (KV70), in both cases from single representative biological replicates (see Materials and methods for detailed explanation). (**E**) GFP-RecN trajectories from representative *ΔrecN ΔrecA* pARA cells (KV71) showing the distance of GFP-RecN foci to midcell relative to cell length. Each shade of green indicates a tracked focus. Midcell distance illustration created in BioRender (https://BioRender.com/y04v494). (**F**) Percentage of GFP-RecN foci colocalizing with DNA versus time after CIP exposure for *ΔrecN ΔrecA* pARA and pSOS cells (KV71 and KV70). (**G**) Mean number of GFP-RecN foci per cell versus time after CIP exposure for *ΔrecN ΔrecA* pARA and pSOS cells (KV71 and KV70). Lines represent means from three biological replicates, and shaded regions indicate SEM.

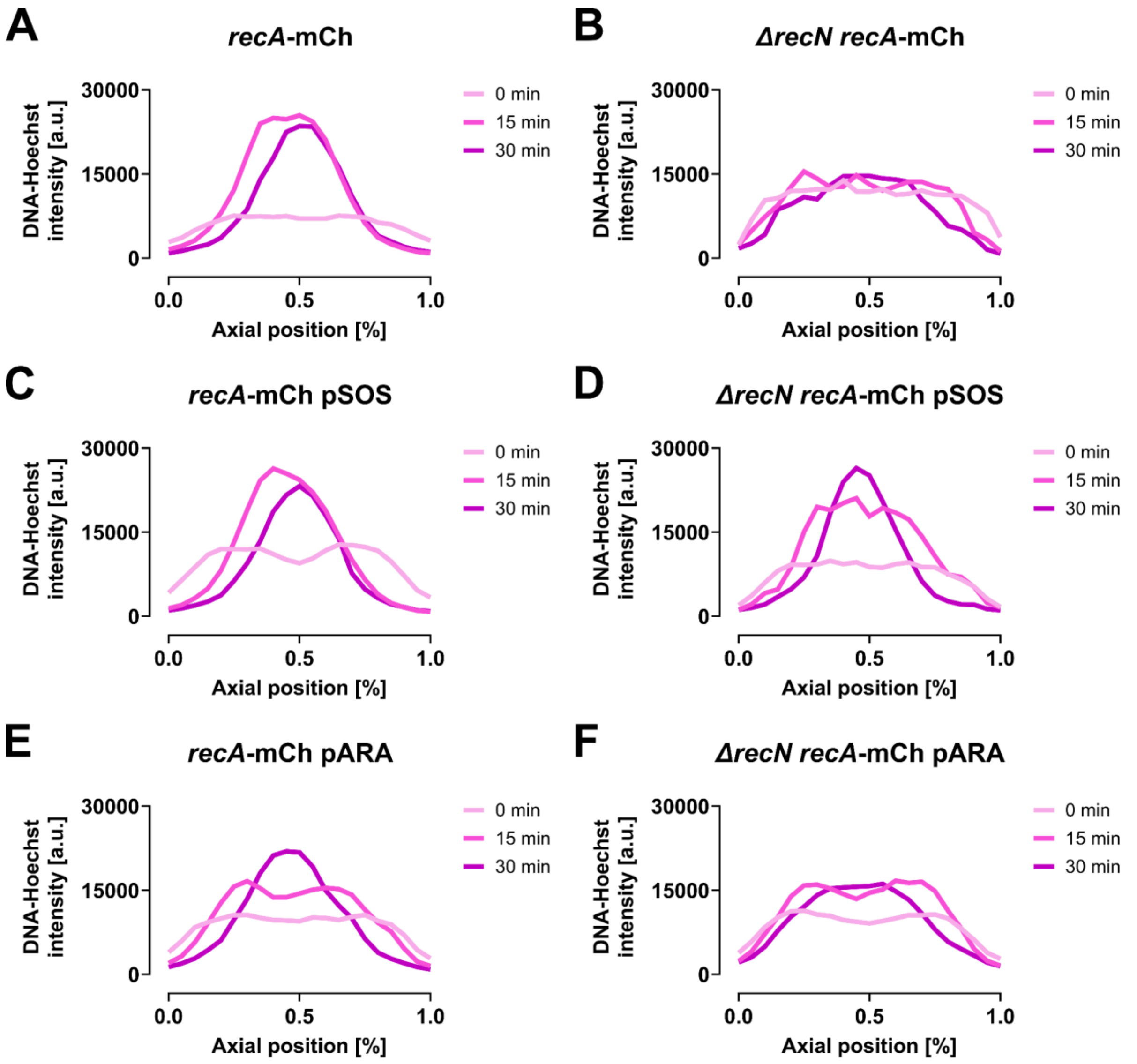


**Figure S6.** Analysis of DNA distribution inside all strains with chromosomally expressed RecA-mCherry to evaluate if this expression alone or in combination with GFP-RecN affects DNA supercompaction. All cells were grown in LB at 37°C, expressing RecA-mCherry chromosomally alongside native RecA. The cells were fixed before, and after 15 and 30 minutes of CIP exposure (10 µg/mL), and then stained with Hoechst 33258. Average Hoechst fluorescence intensity along the cells' long axis for each time point for (**A**) wild type (KV75) from 72-93 cells, (**B**) *ΔrecN* (KV76) from 23-27 cells, (**C**) wild type pARA (KV79) from 31-69 cells, (**D**) *ΔrecN* pARA (KV80) from 46-70 cells, (**E**) wild type pSOS (KV77) from 42-80 cells, and (**F**) *ΔrecN* pSOS (KV78) from 34-41 cells. Results are from single biological replicates (see Materials and methods for detailed explanation). A.u., arbitrary unit.

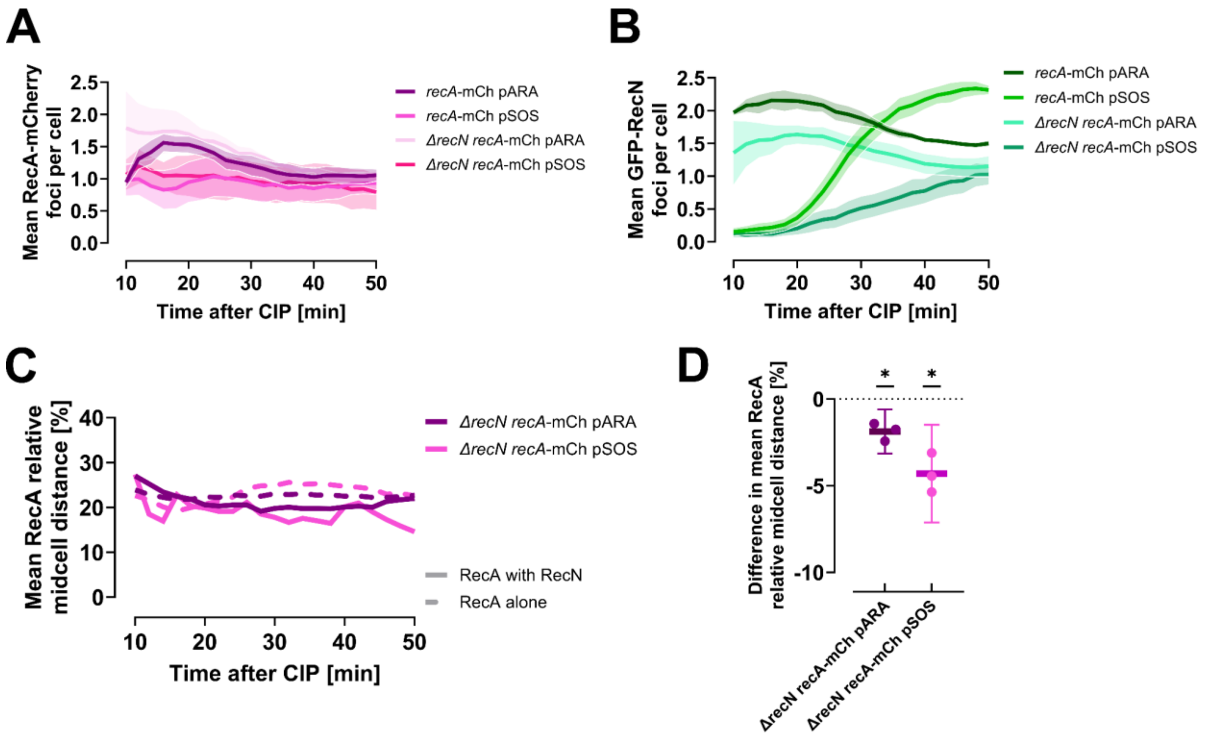


**Figure S7.** Foci count over time for RecA-mCherry and GFP-RecN foci in wild-type and *ΔrecN* strains, and analysis of RecA-mCherry midcell distance in *ΔrecN* background. All cells were grown in LB at 37°C, expressing RecA-mCherry chromosomally alongside native RecA. Cells were imaged at 2-minute intervals starting 10 minutes post-CIP exposure using live-cell spinning disk microscopy. Results represent three biological replicates. (**A**) Mean number of RecA-mCherry foci per cell versus time after CIP exposure for wild-type and ΔrecN cells with GFP-RecN expression from pARA or pSOS. Lines represent means from three biological replicates, and shaded regions indicate SEM. (**B**) Mean number of GFP-RecN foci per cell versus time after CIP exposure of wild-type and ΔrecN cells with GFP-RecN expression from pARA or pSOS. Lines represent means from three biological replicates, and shaded regions indicate SEM. (**C**) Mean distance of colocalizing (solid line) and non-colocalizing (dashed line) RecA-mCherry foci to midcell relative to cell length in *ΔrecN* cells. Means are from three biological replicates. (**D**) Difference in relative midcell distance for colocalizing versus non-colocalizing RecA-mCherry foci in *ΔrecN* cells. Negative values indicate that RecA-mCherry foci colocalizing with GFP-RecN are closer to midcell than other RecA-mCherry foci. Lines show means, dots represent replicates' mean values, and error bars indicate 95% CIs; * $p \leq 0.05$.

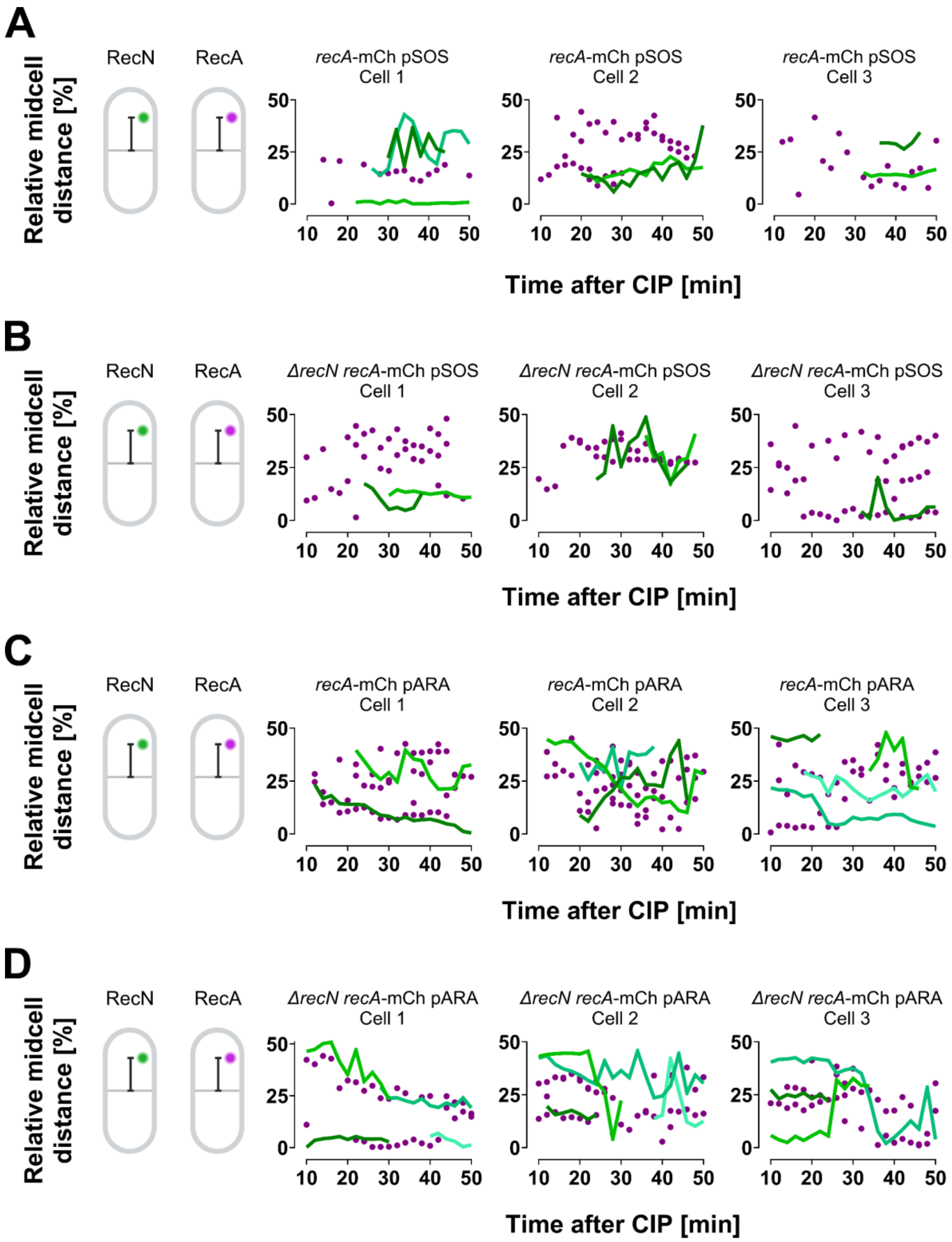


**Figure S8.** Examples of GFP-RecN trajectories and RecA-mCherry localization for representative cells from strains with chromosomally expressed RecA-mCherry. (**A**) Wild type pARA (KV79), (**B**) *ΔrecN* pARA (KV80), (**C**) wild type pSOS (KV77), and (**D**) *ΔrecN* pSOS (KV78). Plots display the distance of foci to midcell relative to cell length, with GFP-RecN trajectories in shades of green and RecA-mCherry foci as magenta dots. Midcell distance illustrations created in BioRender (https://BioRender.com/y04v494).

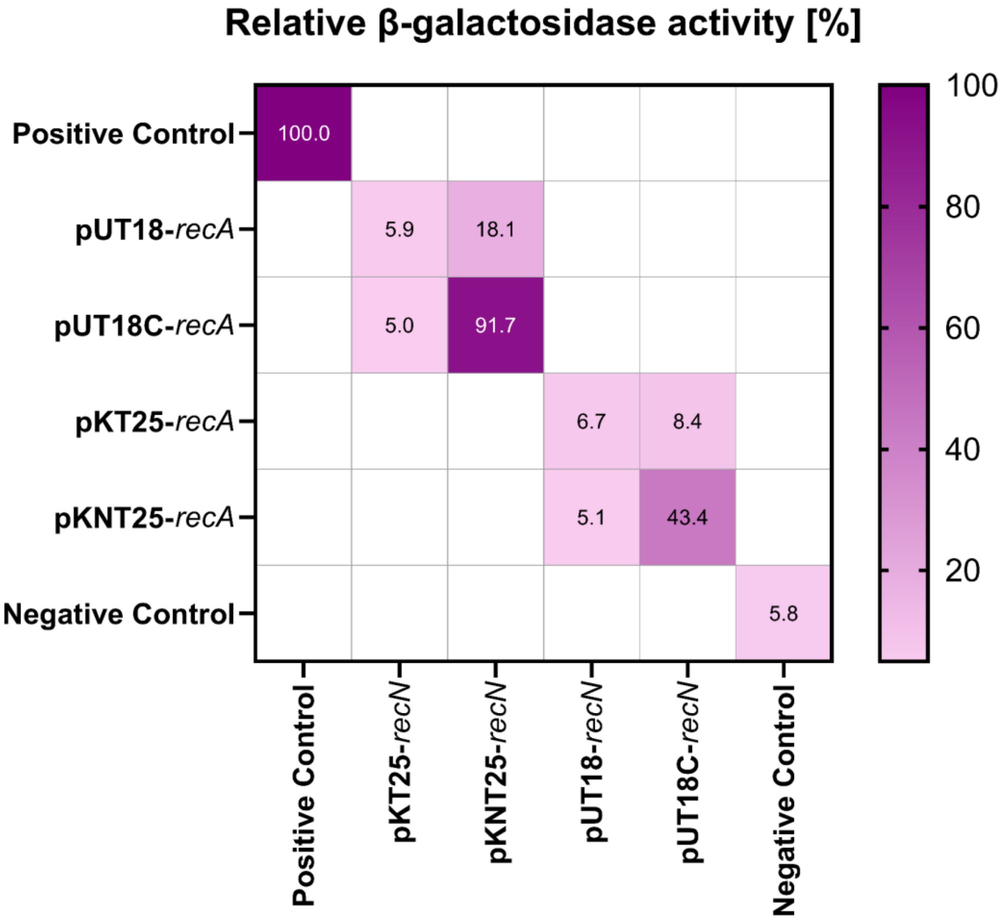


**Figure S9**. Relative β-galactosidase activity results from bacterial two hybrid assay of RecN and RecA interaction. Presented results are means from three biological replicates, each biological replicate averaged from three technical replicates. Relative β-galactosidase activity was calculated versus the positive control of each biological replicate.

**Table S2.** Relative β-galactosidase activity results from bacterial two hybrid assay of RecN and RecA interaction. Presented results are means from three biological replicates, each biological replicate averaged from three technical replicates. Relative β-galactosidase activity was calculated versus the positive control of each biological replicate. Pos, Positive control; Neg, Negative control. Mean ± SEM.

| **RecN plasmid** | **RecA plasmid** | **Relative β-galactosidase activity [%]** |
|---|---|---|
| pKT25-zip (Pos) | pUT18C-zip (Pos) | 100.0 ± 0.0 |
| pKT25 (Neg) | pUT18C (Neg) | 5.8 ± 3.1 |
| pKT25-*recN* | pUT18-*recA* | 5.9 ± 1.2 |
| pKT25-*recN* | pUT18C-*recA* | 5.0 ± 0.9 |
| pKNT25-*recN* | pUT18-*recA* | 18.1 ± 12.3 |
| pKNT25-*recN* | pUT18C-*recA* | 91.7 ± 25.1 |
| pUT18-*recN* | pKT25-*recA* | 6.7 ± 0.4 |
| pUT18-*recN* | pKNT25-*recA* | 5.1 ± 1.7 |
| pUT18C-*recN* | pKT25-*recA* | 8.4 ± 2.3 |
| pUT18C-*recN* | pKNT25-*recA* | 43.4 ± 22.5 |

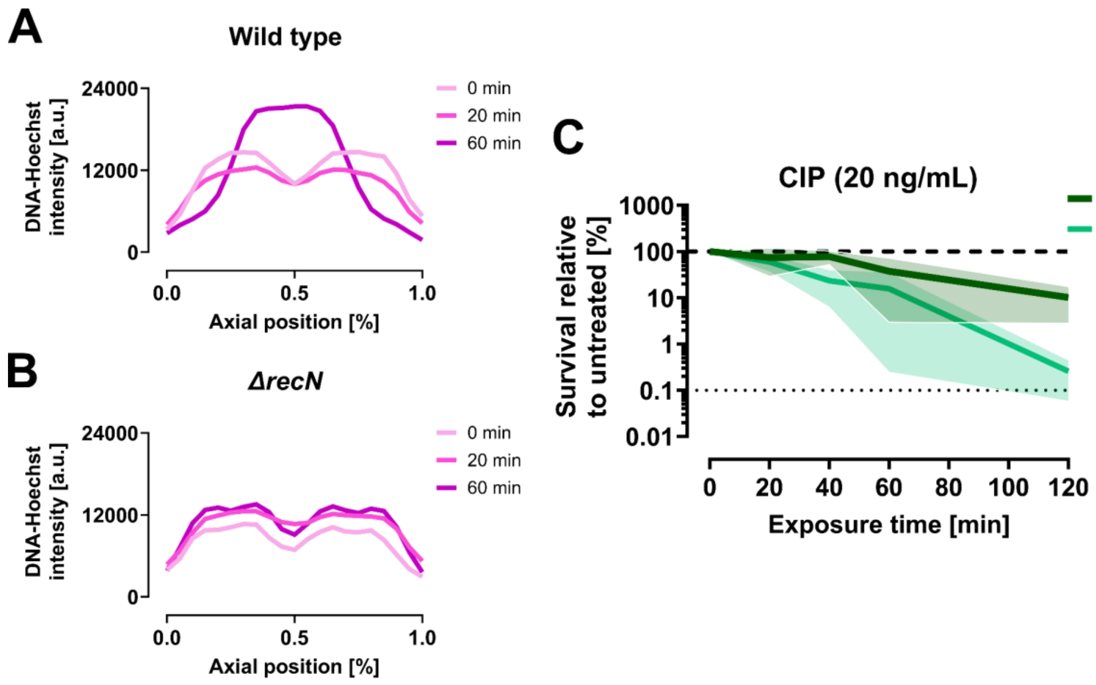


**Figure S10.** *ΔrecN* cells are overly sensitive to CIP also at the MIC dose and fail to compact DNA like wild-type cells within 60 minutes after exposure to MIC doses of CIP. (**A** and **B**) Average Hoechst fluorescence intensity along the cells' long axis for (**A**) wild-type (BW25113) or (**B**) *ΔrecN* (JW5416) cells fixed before or after 20 or 60 minutes of CIP exposure at MIC dose (20 ng/mL) and stained with Hoechst 33258. Results for each time point are averaged from 35-137 cells from a single biological replicate (see Materials and methods for detailed explanation). A.u., arbitrary unit. (**C**) Time-dependent survival assay for wild-type (BW25113) and *ΔrecN* (JW5416) cells after exposure to 20 ng/mL CIP (MIC for wild-type cells). Exposure time refers to the duration of CIP exposure before washing and plating the cells. Survival was measured as colony forming units (CFU) per mL and relative survival was calculated by comparing with untreated parallels. Thick dashed lines indicate survival of untreated parallel; thin dotted lines indicate the assay's detection limit. Green lines represent means of 3-4 biological replicates; shaded regions indicate standard deviation.
